# Epitope-based chimeric peptide vaccine design against S, M and E proteins of SARS-CoV-2 etiologic agent of global pandemic COVID-19: an *in silico* approach

**DOI:** 10.1101/2020.03.30.015164

**Authors:** M. Shaminur Rahman, M. Nazmul Hoque, M. Rafiul Islam, Salma Akter, A. S. M. Rubayet-Ul-Alam, Mohammad Anwar Siddique, Otun Saha, Md. Mizanur Rahaman, Munawar Sultana, M. Anwar Hossain

## Abstract

Severe acute respiratory syndrome coronavirus 2 (SARS-CoV-2) is the cause of the ongoing pandemic of coronavirus disease 2019 (COVID-19), a public health emergency of international concern declared by the World Health Organization (WHO). An immuno-informatics approach along with comparative genomic was applied to design a multi-epitope-based peptide vaccine against SARS-CoV-2 combining the antigenic epitopes of the S, M and E proteins. The tertiary structure was predicted, refined and validated using advanced bioinformatics tools. The candidate vaccine showed an average of ≥ 90.0% world population coverage for different ethnic groups. Molecular docking of the chimeric vaccine peptide with the immune receptors (TLR3 and TLR4) predicted efficient binding. Immune simulation predicted significant primary immune response with increased IgM and secondary immune response with high levels of both IgG1 and IgG2. It also increased the proliferation of T-helper cells and cytotoxic T-cells along with the increased INF-γ and IL-2 cytokines. The codon optimization and mRNA secondary structure prediction revealed the chimera is suitable for high-level expression and cloning. Overall, the constructed recombinant chimeric vaccine candidate demonstrated significant potential and can be considered for clinical validation to fight against this global threat, COVID-19.

## Introduction

Emergence of novel coronavirus, known as severe acute respiratory syndrome coronavirus 2 (SARS-CoV-2), which was first reported in Wuhan, Hubei Province, China in December 2019 is responsible for the ongoing global pandemic of coronavirus disease 2019 (COVID-19). The ongoing outbreak of COVID-19 has already made its alarmingly quick transmission across the globe in 199 countries and regions with a total of 723,077 active infection cases and 33,983 deaths until March 29, 2020^1^. World Health Organization (WHO) has declared a global health emergency, and scientists are racing over the clock to develop effective interventions for controlling and preventing COVID-19 by analyzing novel SARS-CoV-2 genomics, functional structures, and its host-pathogen interactions. Efforts to contain the virus are ongoing; however, given the many uncertainties regarding pathogen transmissibility and virulence, the effectiveness of these efforts is unknown.

This SARS-CoV-2 is the third coronovairus (CoV) that can infect human after the two previously reported coronavirus-severe acute respiratory syndrome (SARS-CoV)^2, 3^ and Middle East respiratory syndrome (MERS-CoV)^4–6^. Alike SARS-CoV and MERS-CoV, the recent SARS-CoV-2 is a positive-sense single-stranded RNA virus (+ssRNA) belonging to Genus *Betacoronavirus* within the family Coronaviridae^7^. The genome of SARS-CoV-2 is approximately 30 kilobases (between 26,000 and 32,000 bases) in length^8^, and encodes for multiple structural and non-structural proteins^7^. The four major structural proteins of CoVs are the spike (S) glycoprotein, small envelope protein (E), membrane protein (M), and nucleocapsid protein (N)^7^. Among these proteins, the S glycoprotein is composed of two main domains, the receptor-binding domain (RBD) and N-terminal domain (NTD),^9, 10^ which play essential role for the attachment and entry of viral particles into the host cells^11, 12^ and characterized by high antigenicity and surface exposure^2, 11, 13^.

To combat the devastating effects of SARS-CoV-2, Scientists applied different vaccine strategies globally. In addition, a curative approach using different drug candidates have been tried so far to combat the current pandemic COVID-19. However, COVID-19 is marching too fast and reached almost every territory in earth. Hence, apart from developing novel drugs against this viral infection, a very potent vaccine candidate is very much desirable. Being an RNA virus, the virus had undergone mutations several times already in the last 4 months^14^. As a result, instead of single epitope, multi-epitope based vaccines can play a great role in fighting against this novel CoV.

On coronaviruses, the component primarily targeted by monoclonal antibodies (mAbs) is the trimeric S glycoprotein of the virion consisting of three homologous chains (A, B and C), which mediate receptor recognition and membrane fusion during viral entry into sensitive cells through the host angiotensin-converting enzyme 2 (ACE2) receptor^5, 9, 11, 13^. However, with the rapid evolution and high genomic variation of RNA viruses, mutations on the RBD may enable the new strains to escape neutralization by currently known RBD-targeting antibodies. Therefore, other functional regions of the S glycoprotein can be important for developing effective prophylactic and therapeutic interventions against SARS-CoV-2 infection. Several mAbs targeting non-RBD regions, especially the NTD has recently been reported^13, 15^. The NTD is located on the side of the spike trimer and has not been observed to undergo any dynamic conformational changes^11^, and therefore, might play role in viral attachment, inducing neutralizing antibody responses and stimulates a protective cellular immunity against SARS-CoV^2, 6, 11^. Besides, administration of RBD and NTD proteins also induced highly potent neutralizing antibodies and long-term protective immunity in animal models^9^. Furthermore, the E and M proteins also have important functions in the viral assembly of a coronavirus and can augment the immune response against SARS-CoV^11, 16, 17^. Monoclonal antibodies with potent neutralizing activity have become promising candidates for both prophylactic and therapeutic interventions against SARS-CoV-2 infections^13^. Therefore, the generation of antibodies targeting the RBD and/or NTD of the S glycoprotein, M and E proteins of SARS-CoV-2 would be an important preventive and treatment strategy that can be tested further in suitable models before clinical trials^18^. The nucleocapsid (N) protein of SARS-coronavirus (SARS-CoV), buried inside phospholipid bilayer, is the major protein in the helical nucleocapsid of virion. This protein is reported to be more conserved than other structural proteins, an important B cell immunogen, as well as, can elicit long lasting and persistent cellular immune responses. Nevertheless, we did not consider this protein in the chimeric vaccine formation because of its initial unavailability outside of the host cell during infection^19^.

Most previous studies of the CoVs have focused on the use of single recombinant antigens, however, the immune responses generated have been inadequate to warrant their use in the development of an effectual protective tool^20, 21^. Alternatively, we propose the development of a multi-epitope vaccine candidate which could lead to the generation of a more potent protective immune response. Multi-epitope vaccine candidates have already been designed for several diseases, including FMD (in our laboratory; unpublished data), MERS and SARS, and their efficacies have been further reported in MERS-CoV and SARS-CoV^2, 6, 22^. Hence, we can assume that chimeric vaccine targeting multiple epitopes on the RBD and NTD segments of the S protein, M and E proteins would be a potentially effective vaccine candidate in combatting SARS-CoV-2, and therefore, could be used for vaccine design. Combining subunit vaccines with established or new adjuvants may represent a faster and safer strategy to move through early clinical development and analyze the efficacy. This study was therefore aimed to *in-silico* design and construction of a multi-epitope-based chimeric vaccine candidate against the globally pandemic highly pathogenic SARS-CoV-2.

## Results

### Retrieving protein sequences

We retrieved the spike (S), envelop (E) and membrane (M) protein sequences of the SARS-CoV-2, and the S protein sequences of the SARS-CoV and MERS-CoV from the whole genome reference sequences of the respective three viruses from the NCBI database. To map the structural variability of the S protein of SARS-CoV-2, SARS-CoV and MERS-CoV, we applied homology search on the respective S protein sequences, and finally selected three protein data bank (PDB) templates such as 6vsb, 6acd and 5w9h, respectively for predicting the three dimensional (3D) conformation of the S proteins. Finally, the 3D conformation of the S proteins of the respective three viruses were validated using the Ramachandran plot analysis (Supplementary Fig. 1).

### Physicochemical properties and trimeric structure of spike (S) protein

The physiochemical properties of the spike (S) glycoproteins of the SARS-CoV-2, SARS-CoV and MERS-CoV computed using the Protparam on the ExPASy server demonstrates that it varies significantly among the three viruses. The theoretical isoelectric points (PI) of the subject proteins were 6.24, 5.56 and 5.70, respectively. Protparam computed instability-index (II) of the S protein of SARS-CoV-2, SARS-CoV and MERS-CoV were 33.01, 32.42 and 36.60 indicating that these proteins are quite stable^4, 23^.

The S protein of the SARS-CoV-2, SARS-CoV and MERS-CoV is a trimeric conformation consisting of three homologous chains naming as chain A, B and C^5, 10^ (Fig. 1). The individual structural domain of coronavirus S proteins reflects high degree of structural divergences in the receptor-binding domain (RBD) and N-terminal domain (NTD) of the chain A and chain C compared to that of chain B. Multiple sequence alignment revealed that the S protein of SARS-CoV-2 shares 77.38% and 31.93% sequence identity with the S proteins of the SARS-CoV and MERS-CoV, respectively, and these findings are in line with many earlier studies^3–7^ (Supplementary Fig. 2). Due to structural divergence in RBD and NTD of S proteins in SARS-CoV-2, SARS-CoV and MERS-CoV, these regions are the main focus of structural studies including epitope-based chimeric vaccine candidate designing, selection, and development.

**Fig. 1.**
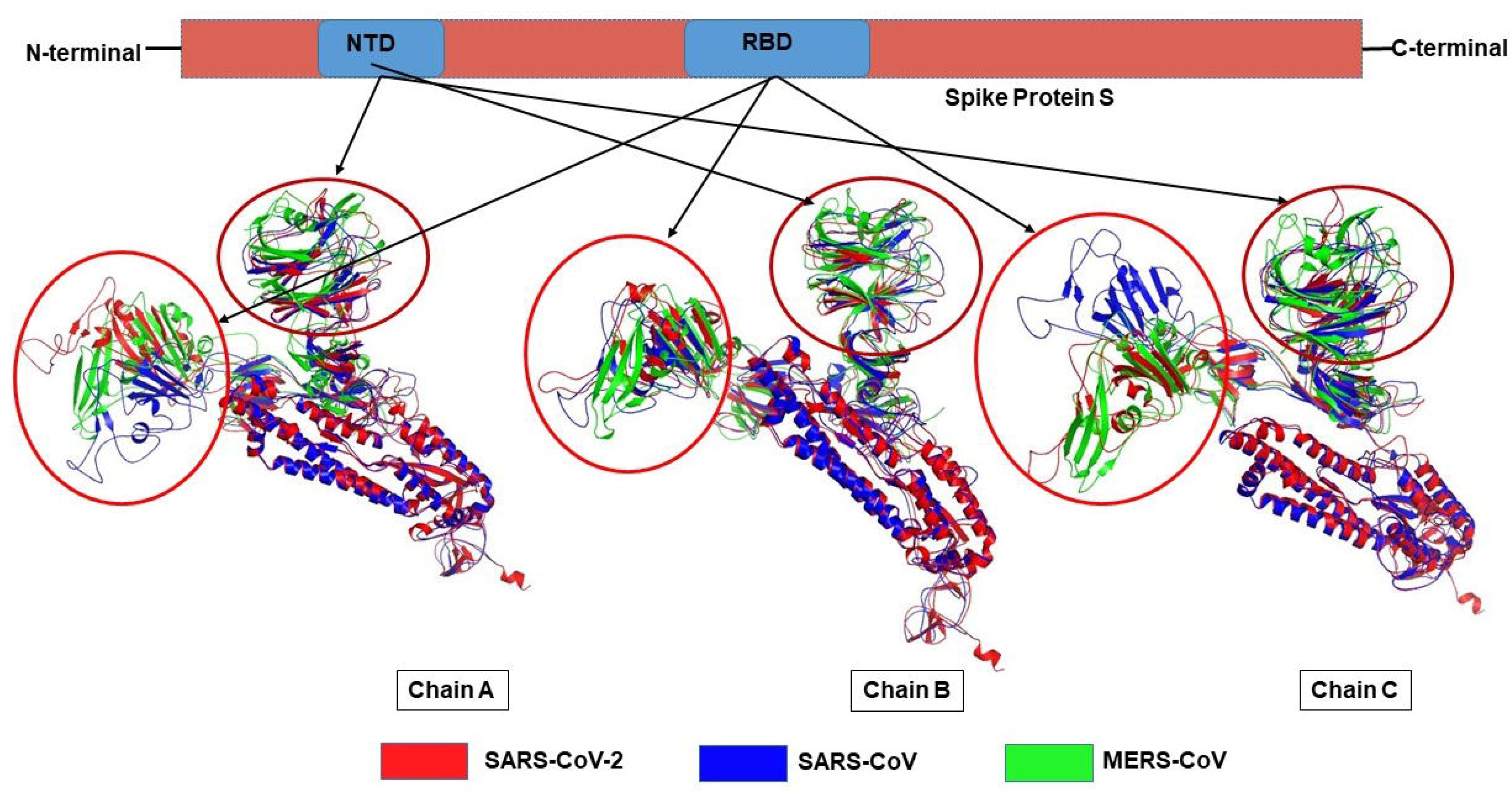
The trimeric conformation spike (S) proteins. The S proteins of the SARS-CoV-2, SARS-CoV and MERS-CoV is a trimeric conformation consisting of three homologous chains named as chain A, B and C. Respective sequences of these three chains were aligned and visualized using PyMOL which revealed high degree of structural divergences in the N-terminal domains (NTDs) and receptor binding domains (RBDs) of the chain A and C compared to that of chain B.

### Structure-based B-cell epitopes prediction

Linear epitopes prediction based on solvent-accessibility and flexibility revealed that chain A of S protein possessed 15 epitopes, of which three epitopes having different residues positions (395-514, 58-194, 1067-1146) were highly antigenic (score > 0.8). However, the B and C chains had 18 and 19 epitopes, respectively, and of them three epitopes in chain B (residues position 1067-1146, 89-194, 58-87) and two epitopes in chain C (residues position 56-194, 1067-1146) seem to be highly antigenic (score > 0.8) (Table 1). The amino acid residues 395-514 and 56-194 of the detected epitopes belonged to RBD and NTD regions of the S protein, respectively. These regions were considered as the potential epitope candidates in the IEDB analysis resource Elipro analysis. However, the epitopes with amino-acids position of 1067-1146 were not selected as the potential epitope candidate because of their presence in viral transmembrane domains (Supplementary Fig. 3). The tertiary structures of the RBD and NTD illustrate their surface-exposed position on the S protein (Fig. 2).

**Fig. 2.**
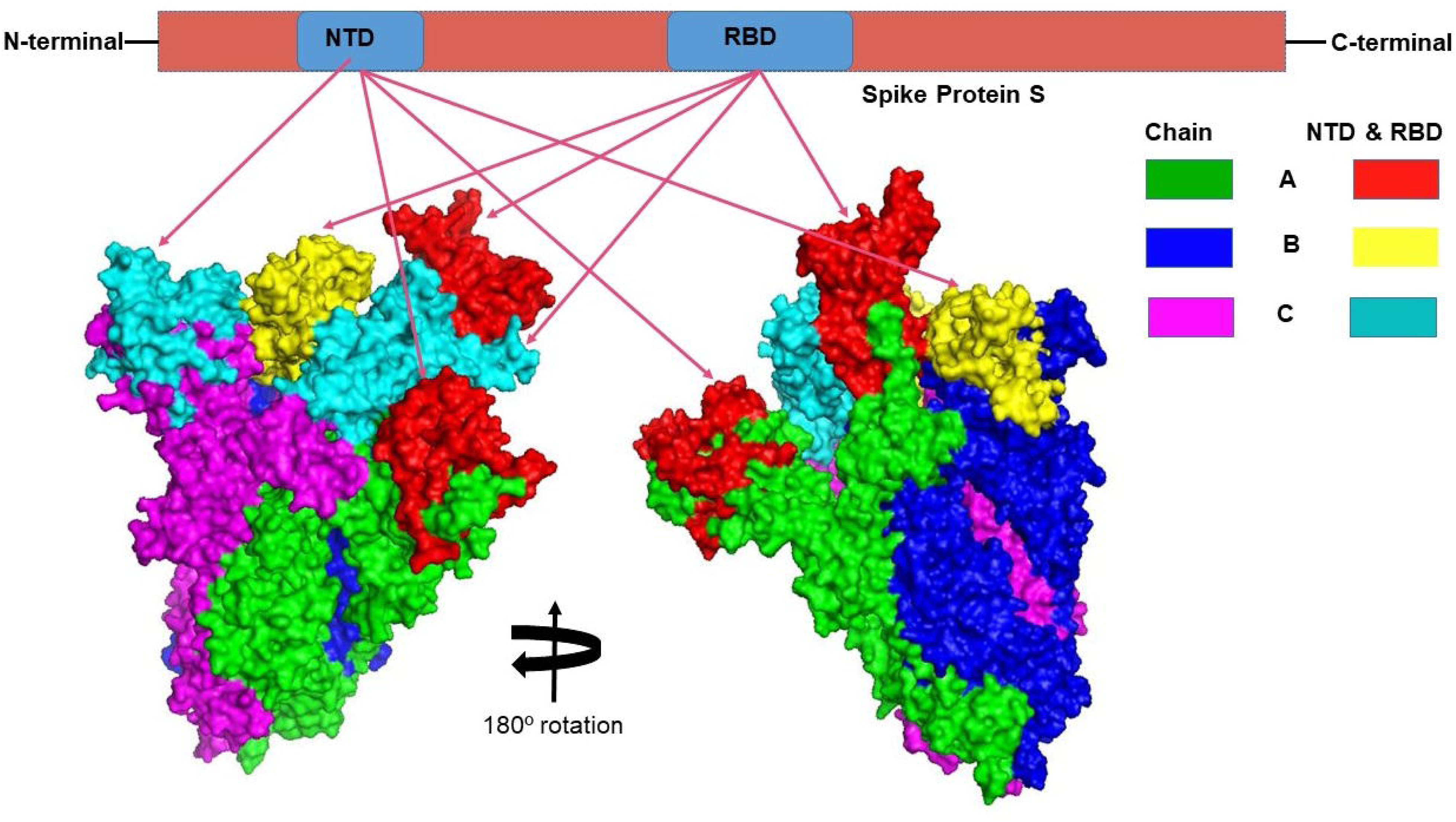
The three-dimensional (3D) structure of the N-terminal domains (NTDs) and receptor binding domains (RBDs) of the spike (S) proteins of SARS-CoV-2 (surface view). The red, cyan, and yellow colored regions represent the potential antigenic domains predicted by the IEDB analysis resource Elipro analysis.

**Table 1:**
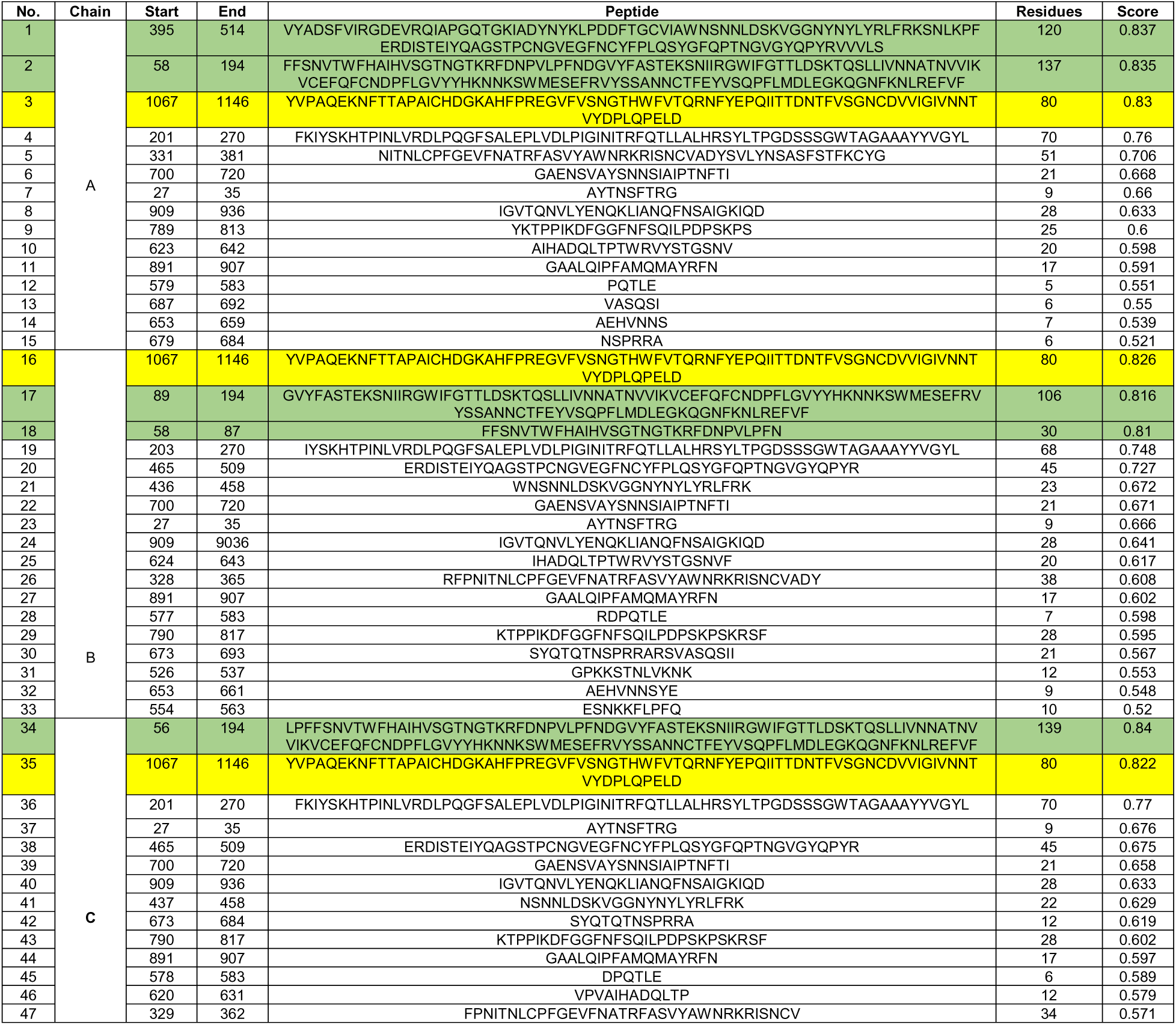
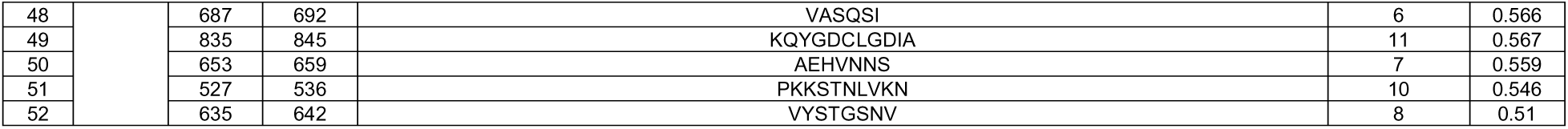
Linear epitopes present on spike (S) glycoprotein surface predicted through ElliPro in IEDB-analysis resource based upon solvent-accessibility and flexibility are shown with their antigenicity scores. The highlighted green coloured regions were the potential antigenic domains while the yellow coloured region represents the trans-membrane domain of the S protein.

### Sequence-based B-cell epitopes

Antigenicity analysis of full-length S glycoprotein, M (membrane) and E (envelope) proteins using VaxiJen v2.0 showed antigenicity scores of 0.465, 0.510 and 0.603, respectively exhibiting them as potential antigens. Therefore, they were considered to be promising epitopes for vaccine candidate selection against globally pandemic SARS-CoV-2 infections. A total of 22 B-cell epitopes were predicted in S protein of the SARS-CoV-2 using IEDB analysis resource and Bepipred linear epitope prediction 2.0 tools. Of the detected epitopes, eight and six epitopes were respectively found in RBD (aa position 328-528) and NTD (aa position 56-194) regions while the E and M proteins had 2 and 6 epitopes, respectively (Fig. 3, Supplementary Table 1). However, only 5 epitopes were exposed on the surface of the virion, and had a high antigenicity score (> 0.4), indicating their potentials in initiating immune responses (Table 2). To find out the most probable peptide-based epitopes with better confidence level, selected peptides were further tested using VaxiJen 2.0 tool, and five peptides having antigenicity score of ≥ 0.5 were annotated. Among the 5 annotated epitopes, RBD region had two highly antigenic epitopes (75-IRGDEVRQIAPGQTGKIADYNYKLPD-100, antigenicity score = 0.932 and 166-QSYGFQPTNGVGYQ-189, antigenicity score = 0.668). Similarly, the NTD region harbored two highly antigenic epitopes (117-SQPFLMDLEGKQGNFKNLRE-136, antigenicity score = 0.749 and 17-GTNGTKRFDN-26, antigenicity score = 0.667). In addition, the envelop (E) protein possessed only one highly antigenic epitope (57-YVYSRVKNLNSSRVP-71, antigenicity score = 0.449) which might be potentially functional in host cell binding (Table 2). Furthermore, the Kolaskar and Tongaonkar antigenicity profiling found five highly antigenic epitopes in RBD region with an average (antigenicity) score of 1.042 (minimum = 0.907, maximum = 1.214), and seven highly antigenic epitopes in NTD with an average (antigenicity) score of 1.023 (minimum = 0.866, maximum = 1.213) (Supplementary Fig. 4, Supplementary Table 2). The average Kolaskar scores for envelop protein B-cell epitope (EBE) and membrane protein B-cell epitope (MBE) were 0.980 and 1.032, respectively (Supplementary Table 2). However, through ABCPred analysis, we identified 18 and 11 B-cell epitopes in RBD and NTD regions with average antigenicity score of 0.775 and 0.773 in the associated domains, respectively (Supplementary Table 3).

**Fig. 3.**
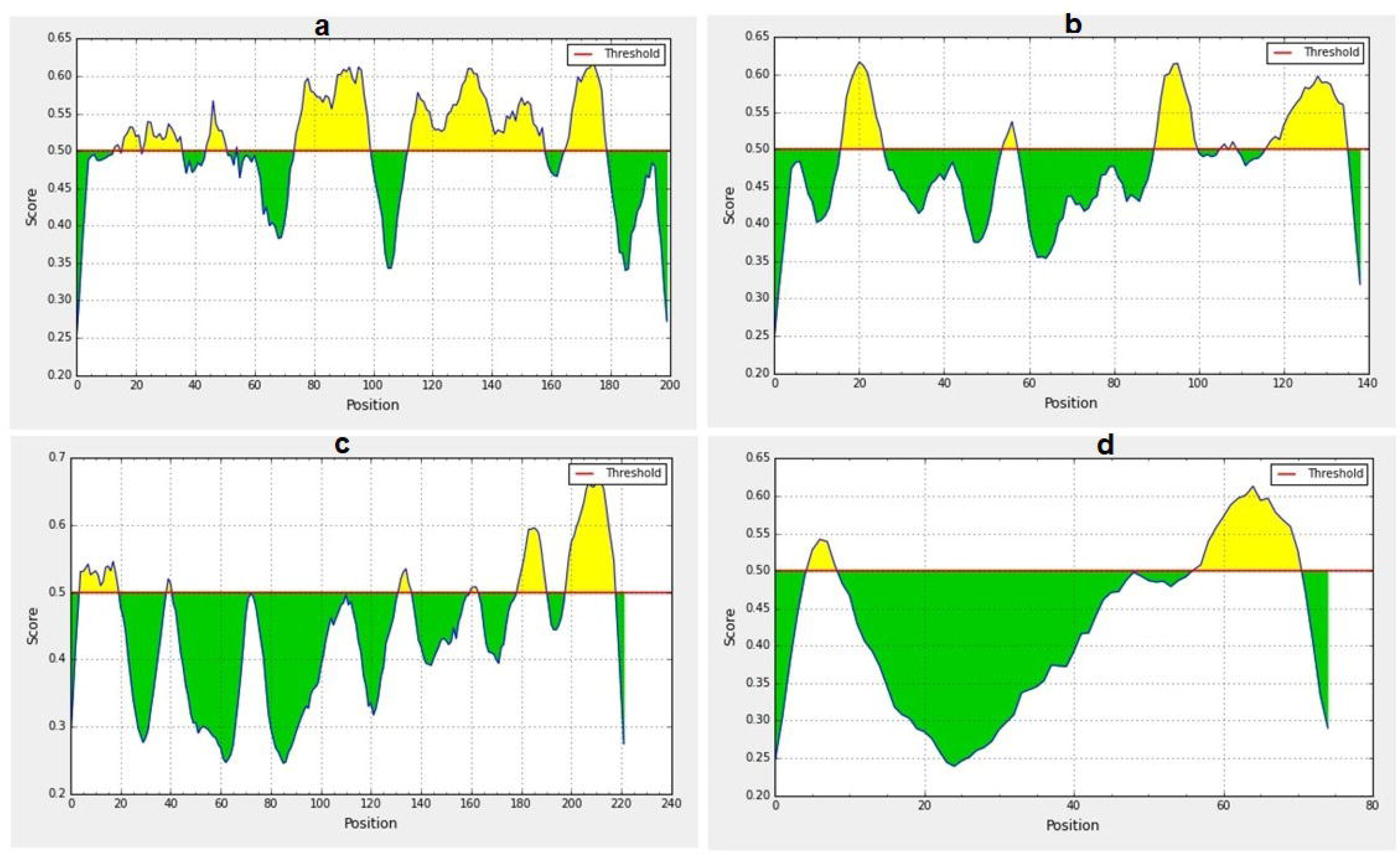
Predicted B-cell epitopes using BepiPred-2.0 epitope predictor in IEDB-analysis resource web-based repository. Yellow areas above threshold (red line) are proposed to be a part of B cell epitopes in (a) RBD and (b) NTD regions of S protein, (c) envelop (E) and (d) membrane (M) proteins of SARS-CoV-2. While green areas are not.

**Table 2:**
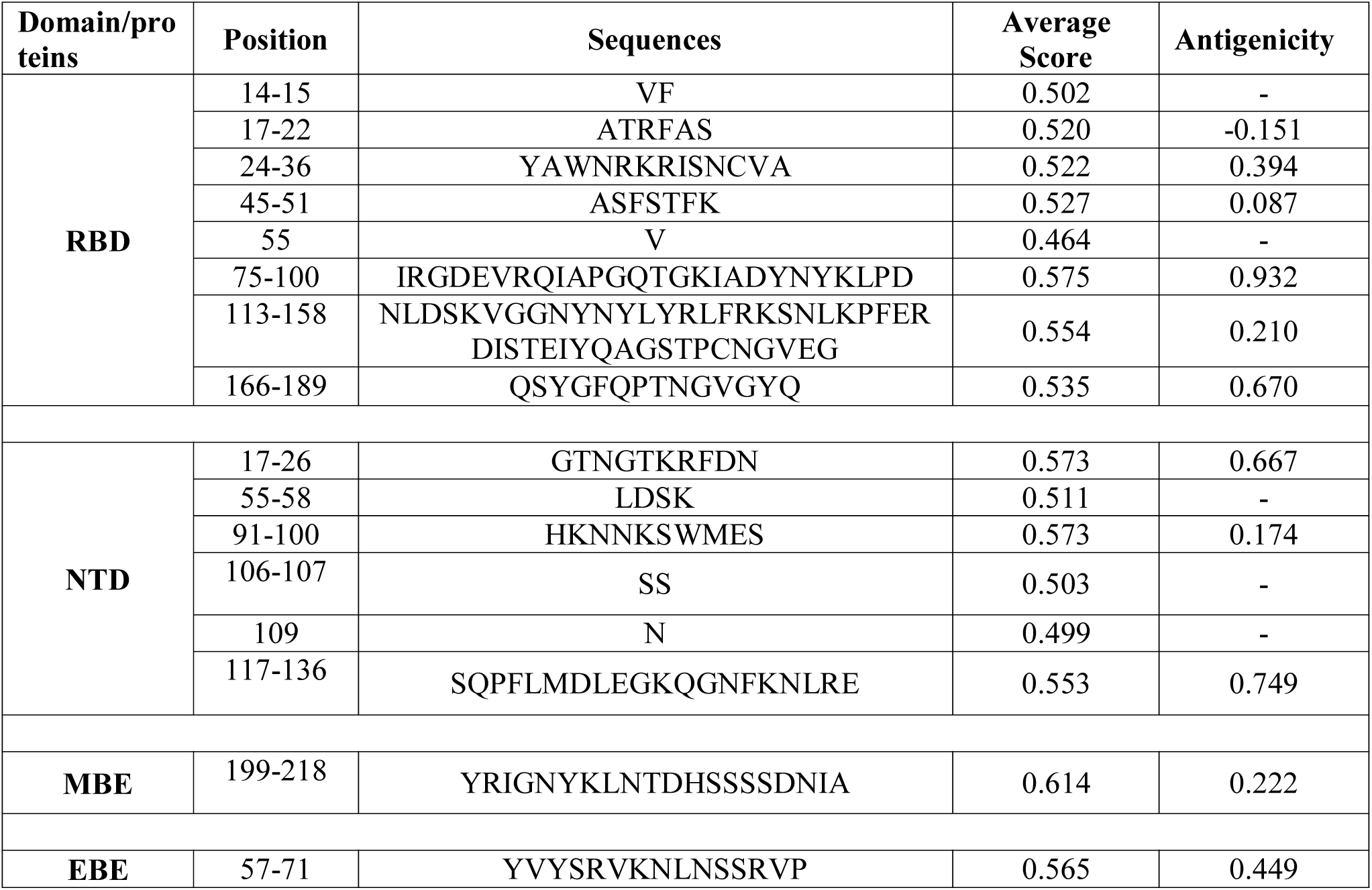
B-cell epitopes predicted using Bepipred linear epitope prediction 2.0 in IEDB analysis resource web-server along with their start and end positions, average score, and VaxiJen 2.0 determined antigenicity scores.

### Recognition of T-cell epitopes

The IEDB analysis resource tool was employed to identify T-cell epitopes in RBD and NTD of the S protein of SARS-CoV-2. The prediction method included the major histocompatibility complex class I and (MHC-I, MHC-II), proteasomal cleavage and TAP transport. The RBD and NTD regions were analyzed using IEDB MHC-I binding prediction tool to predict T lymphocytes epitopes that have binding affinity with MHC-I alleles based on Artificial Neural Network (ANN) with half-maximal inhibitory concentration (IC_50_) ≤ 50 nM. A lower IC_50_ value indicates higher binding affinity of the epitopes with the MHC class I and II molecules. While most of the studies^24, 25^ reported that a binding affinity (IC_50_) threshold of 250 nM identifies peptide binders recognized by T-cells, and this threshold can be used to select peptides, we kept binding affinity within 50 nM to get better confidence level in predicting epitopes for MHC-I and MHC-II alleles. The IEDB MHC-I prediction tool retrieved 77 T-cell epitopes in RBD that interacted with 21 possible MHC-I alleles whereas the NTD domain possessed 35 T-cell epitopes with 17 possible MHC-I alleles (Supplementary Data 1). Similarly, the IEDB MHC-II prediction tool generated 13-mer124 peptides from the RBD, and 10-mer 73 peptides in the NTD segments of the S protein that showed interaction with many different and/or common MHC-II alleles with an IC_50_ value ranging from 1.4 to 49.9 nM (Supplementary Data 1). Furthermore, for MHC-I and MHC-II processing, the analysis tool of the IEDB generates an overall score for each epitope’s intrinsic potential of being a T-cell epitope based on proteasomal processing, TAP transport, and MHC-binding efficiency (Supplementary Data 1). The outcomes of these tools are quite substantial because they utilize vast number of alleles of HLAs (human-leukocyte-antigens) during computation.

### Molecular docking analysis of T-cell epitopes with HLA alleles

From the selected epitopes from the RBD and NTD segments, top five based on IC_50_ score were used in molecular docking analysis using the GalaxyWeb server with their respective HLA allele binders, in which they revealed significantly favorable molecular interaction for binding affinity. Docking complexes thus formed have significantly negative binding energy, and most of the aa residues of the epitopes were involved in molecular interactions with their respective HLA alleles (Supplementary data 1). The epitope-HLA docking complexes were further refined with GalaxyRefineComplex, and their binding affinity was analyzed through PRODIGY web-server. All of the selected epitopes showed significantly negative binding affinity (ΔG always remained ≤ −8.2 kcal mol^-1^, average = −9.94 kcal mol^-1^, Fig. 4, Supplementary data 1).

**Fig. 4.**
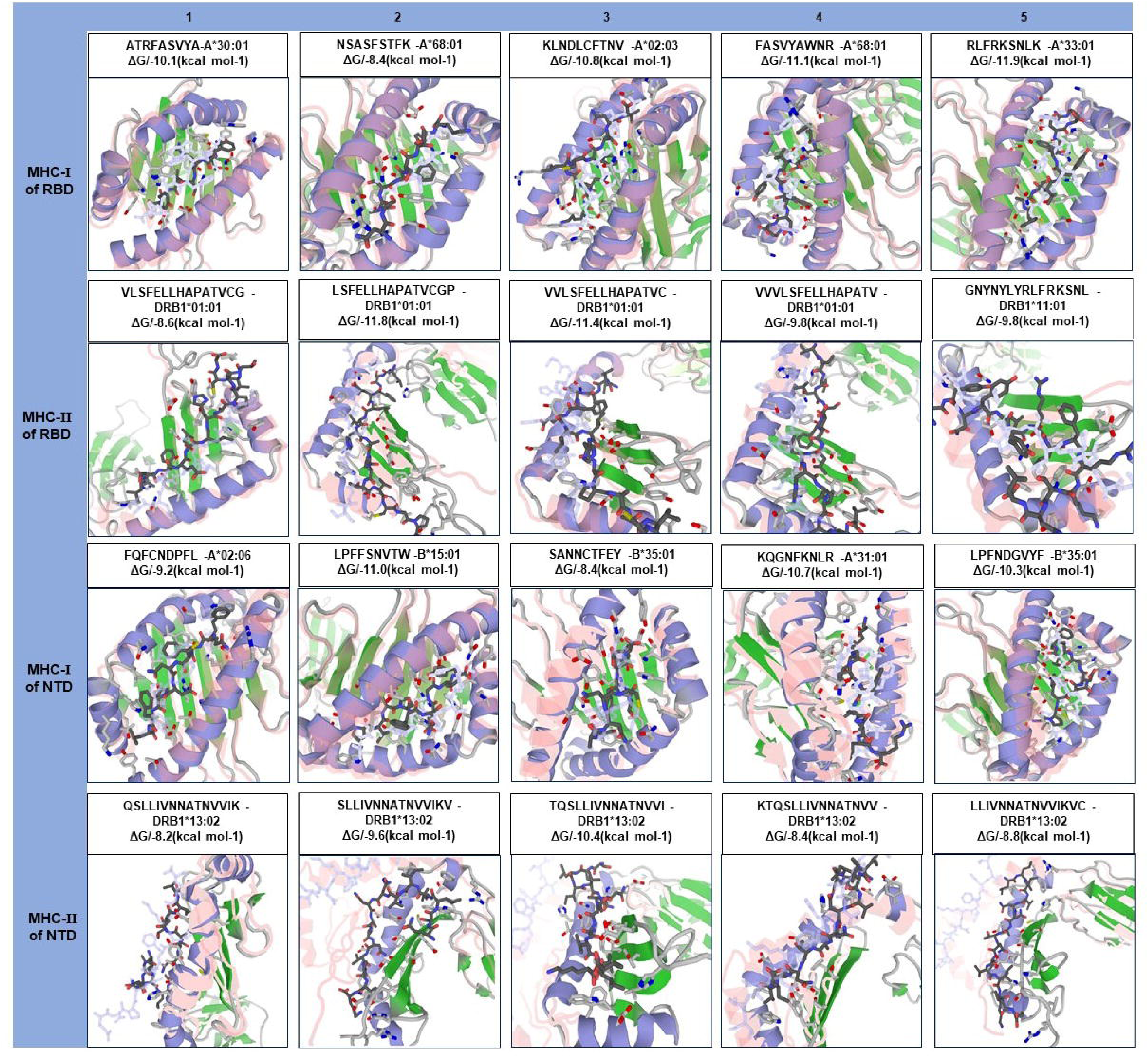
Molecular docking of top five MHC-□ and MHC-□ epitopes of RBD and NTD domains with respect to HLA allele binders. The protein-peptide docking was performed in GalaxyWEB-GalaxyPepDock-server followed by the refinement using GalaxyRefineComplex and free energy (ΔG) of each complex was determined in PRODIGY server. Ribbon structures represent HLA alleles and stick structures represent the respective epitopes. Light color represents the templates to which the alleles and epitopes structures were built. Further information on molecular docking analysis is also available in Supplementary Data 1.

### IFN-**γ** inducing epitope prediction

The findings of IFNepitope program suggests that, both the target RBD and NTD regions of S protein, and B-cell linear epitope (MBE) had great probability to release of IFN-γ with a positive score. A total of 56 potential positive IFN-γ inducing epitopes (15-mer) were predicted for the RBD domain with an average epitope prediction score of 0.255 and the maximum SVM score of 0.625. On the other hand, a total of 33 potential positive epitopes were predicted for the NTD domain with an average epitope prediction score of 0.312 and the maximum SVM score of 0.811. Moreover, the M protein also possessed several IFN-γ inducing epitopes having an average epitope prediction score of 0.980 (Supplementary Table 4).

### Population coverage analysis

The distribution of HLA allele varies between different geographical regions and ethnic groups around the globe. Therefore, population coverage during the development of an efficient vaccine must be taken into account. In this study, epitopes having IC_50_ values ≤ 50 nM and their respective alleles (CTL and HTL) were considered for population coverage analysis both individually (MHC-I and MHC-II) and in combination (Supplementary Data 2). Our selected CTL and HTL epitopes were found to cover 94.9% and 73.11% of the world population, respectively. Importantly, CTL and HTL epitopes showed 98.63% population coverage worldwide when used in combination. The highest population coverage was found to be 99.99% in the Latin American country, Peru (Fig. 5, Supplementary Data 2). In China, where the viral strain (SARS-CoV-2) first appeared and had more devastating outbreaks, the population coverage for CTL and HTL epitopes was 92.67% and 53.44%, respectively with a combined coverage of 96.59%. SARS-CoV-2 is currently causing serious pandemics in different continents of the globe including Italy, England, Spain, Iran, South Korea and United States of America where the combined population coverage was found to be 98.8%, 99.44%, 95.35%, 98.48%, 99.19% and 99.35%, respectively (Fig. 5a, Supplementary Data 2). In addition to geographical distribution, the ethnic groups also found to be an important determinant for good coverage of the CTL and HTL epitopes (Fig. 5b). Of the studied 147 ethnic groups, the Peru Amerindian had highest population coverage for CTL (99.98%) while the HTL epitopes had highest population coverage for Austria Caucasoid (88.44%) (Fig. 5b, Supplementary Data 2). Furthermore, 53.06% of the ethnic groups had a combined population coverage of more than 90.0% for both CTL and HTL epitopes.

**Fig. 5.**
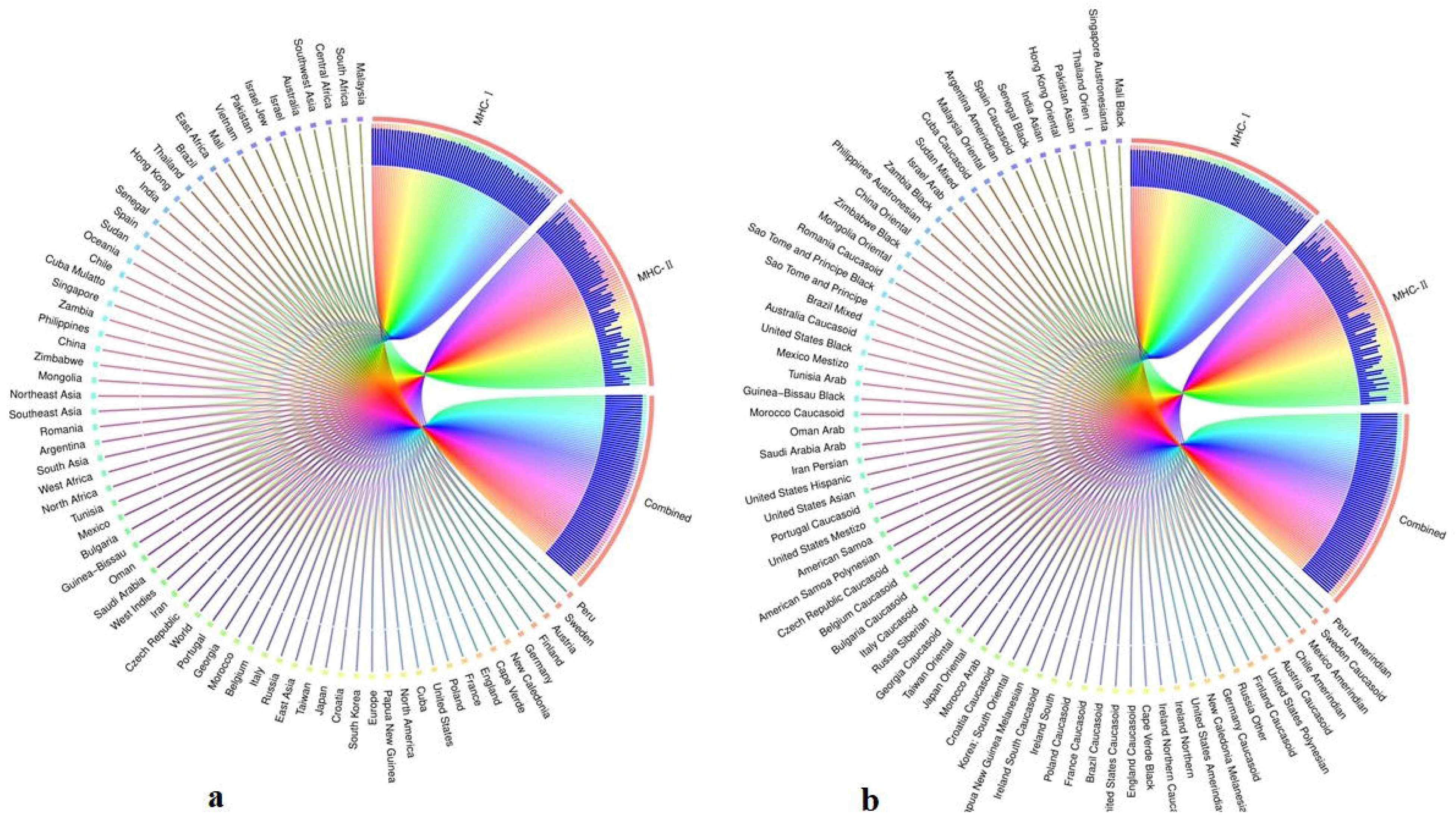
Population coverage of the selected T-cell epitopes and their respective HLA alleles. The circular plot illustrates the relative abundance of the top 70 geographic regions and ethnic groups for selected CTL and HTL epitopes, which were used to construct the vaccine and their corresponding MHC HLA alleles were obtained for population coverage analysis both individually (either MHC-□ or MHC-□) and in combination (MHC-□ and MHC-□). (a) Population coverage of top seventy geographical regions out of 123 regions. (b) Population coverage of top seventy ethnic groups selected from 146 ethnic groups. Regions and ethnic groups in the respective MHC-□ and MHC-□ epitopes are represented by different colored ribbons, and the inner blue bars indicate their respective relative coverages. Further information on population coverage analysis is also available in Supplementary Data 1.

### Design and construction of chimeric vaccine candidate

The selected sequences for designing of chimeric construct were PADRE (13 aa), MBE (20 aa), NTD (139 aa), RBD (200 aa), EBE (15 aa), and Invasin (16 aa), and the construct was named as CoV-RMEN. The schematic diagram of protein domain structures, their junctions and linker sites in CoV-RMEN are shown in Fig. 6a. These fragments were fused together using a repeat of hydrophobic (glycine; G) and acidic aa (glutamic acid; E) linkers. The EGGE, GG, GG, GG and EGGE sequences were introduced between different domains, and epitopes for more flexibility and efficient separation with balancing of acidic and basic amino acids.

**Fig. 6.**
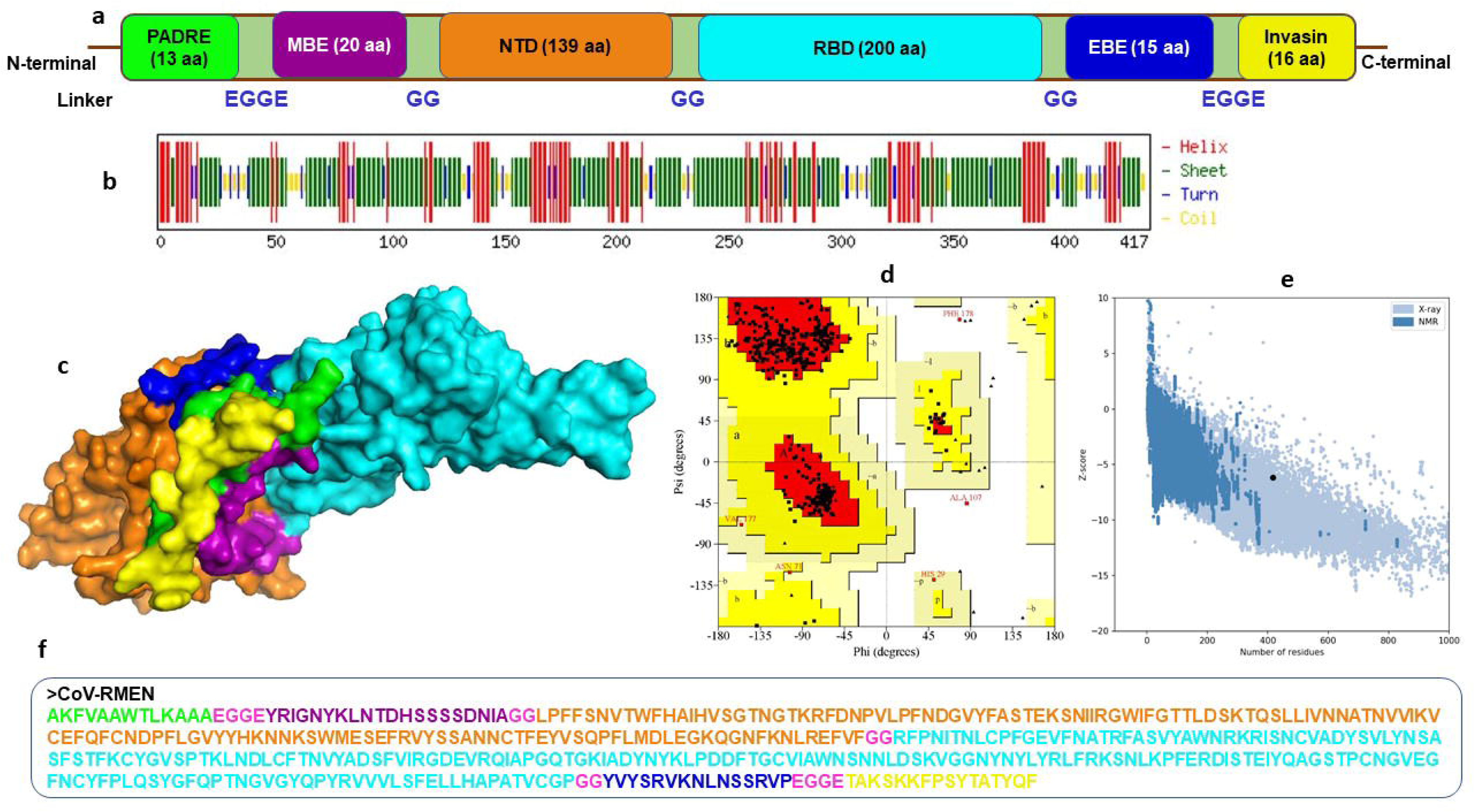
Design, construction and structural validation of multi-epitope vaccine candidate (CoV-RMEN) for SARS-CoV-2. (a) Structural domains and epitopes rearrangement of CoV-RMEN, (b) secondary structure of CoV-RMEN as analyzed through CFSSP:Chou and Fasman secondary structure prediction server, (c) final tertiary structure of CoV-RMEN (surface view) obtained from homology modelling on Phyre2 in which domains and epitopes are represented in different colors (PADRE-green; membrane B-cell epitope, MBE-magenta; N-terminal domain, NTD-orange; receptor-binding domain, RBD-cyan; envelop B-cell epitope, EBE-blue; invasin-yellow), (d) validation of the refined model with Ramachandran plot analysis showing 94.7%, 4.8% and 0.5% of protein residues in favored, allowed, and disallowed (outlier) regions respectively, (e) ProSA-web, giving a Z-score of −6.17, and (f) the finally predicted primary structure of the CoV-RMEN.

### Prediction of the antigenicity and allergenicity of the CoV-RMEN

The antigenicity of the final chimeric protein sequence was predicted by the VaxiJen 2.0 server to be 0.450 with a virus model at a threshold of 0.4 and 0.875 with ANTIGENpro. The results indicate that the generated CoV-RMEN sequences are both antigenic in nature. The vaccine sequence without the adjuvant was also predicted to be non-allergenic on both the AllerTOP v.2 and AllergenFP servers^21^.

### Physiochemical properties and solubility prediction of the CoV-RMEN

The molecular weight (MW) of the finally constructed vaccine candidate was predicted to be 46.8 kDa with a theoretical isoelectric point value (pI) of 8.71. The protein is predicted to be slightly basic in nature based on the pI. The half-life was assessed to be 4.4 hours in mammalian reticulocytes *in vitro*, and >20 hours in yeast and >10 hours in *E. coli in vivo*. The protein was predicted to be less soluble upon expression with a solubility score of 0.330 corroborating the findings of many previous studies^21, 26^. An instability index (II) of 29.74 was predicted, classifying the protein as stable (II of > 40 indicates instability). The estimated aliphatic index was predicted to be 66.59, indicating thermostability. The predicted Grand average of hydropathicity (GRAVY) was −0.300. The negative value indicates the protein is hydrophilic in nature and can interact with water molecules^21^.

### Secondary and tertiary structures prediction of the CoV-RMEN

The CoV-RMEN peptide was predicted to contain 43.2% alpha helix, 67.4% beta sheet, and 12% turns (Fig. 6b, Supplementary Fig. 5) using CFSSP:Chou and Fasman secondary structure prediction server. In addition, with regards to solvent accessibility of aa residues, 34% were predicted to be exposed, 30% medium exposed, and 34% were predicted to be buried. Only 2 aa residues (0.0%) were predicted to be located in disordered domains by the RaptorX Property server (Supplementary Fig. 6). The Phyre2 server predicted the tertiary structure model of the designed chimeric protein in 5 templates (c5x5bB, c2mm4A, c6vsbB, c5x29B and c6vybB) based on heuristics to maximize confidence, percent identity and alignment coverage. The final model of the CoV-RMEN peptide modelled at 82% with more than 90% confidence (Fig. 6c). Moreover, 65 residues were modelled by *ab initio*.

### Tertiary structure refinement and validation of the CoV-RMEN

After energy minimization of the CoV-RMEN using GROMOS96 program^27^, refinement of the vaccine model on the ModRefiner server followed by further refinement on the GalaxyRefine server yielded five models. The selected model has parameters scores of GDT-HA (0.9538), RMSD (0.414), and MolProbity (2.035). The Ramachandran plot analysis of the finally modelled protein revealed that 94.7% of residues in the protein are in favored regions (Fig. 6d), consistent with the 94.0% score predicted by the GalaxyRefine analysis. Additionally, 4.8% of the residues were predicted to be in allowed regions, and only 0.5% in disallowed region (Fig. 6d). The chosen model after refinement had an overall quality factor of 74.45% with ERRAT (Supplementary Fig. 7) while ProSA-web gave a Z-score of −6.17 (Fig. 6e) for the CoV-RMEN protein model.

### Immune simulation

The immune-stimulatory ability of the predicted vaccine CoV-RMEN was conducted through the C-ImmSimm server. The analysis predicts the generation of adaptive immunity in target host species (human) using position-specific scoring matrix (PSSM), and machine learning techniques for the prediction of epitopes and immune interactions^28^. The cumulative results of immune responses after three times antigen exposure with four weeks interval each time revealed that the primary immune response against the antigenic fragments was elevated indicated by gradual increase of IgM level after each antigen exposure (Fig. 7a). Besides, the secondary immune response, crucial for immune stability, have been shown as increased with adequate generation of both IgG1 and IgG2. Also, the elevated level of all circulating immunoglobulins indicates the accuracy of relevant clonal proliferation of B-cell and T-cell population. The level of cytokines after antigen exposure increased concomitantly reflected by escalation of IFN-γ and IL-2 which are most significant cytokines for anti-viral immune response and clonal selection (Fig. 7b). The abundance of different types of B-cells and T-cells, like antigen processing B-cells, resting memory B- and T- cells, B-cells with IgM and IgG remains significantly higher indicating development of immune memory and consequently increased clearance of antigen after exposure (Fig. 7c,d). Additionally, T-helper cells and cytotoxic T-cells were found with a drastic up-regulation of Th1 concentration enhancing the B-cell proliferation and immune memory development (Fig. 7e,f). The high level of immunoglobulin IgG1 + IgG2, active B-cell and T-helper cell population reflected the development of strong immune response reinforcing the indelible and peerless antigenicity of the CoV-RMEN vaccine candidate.

**Fig. 7.**
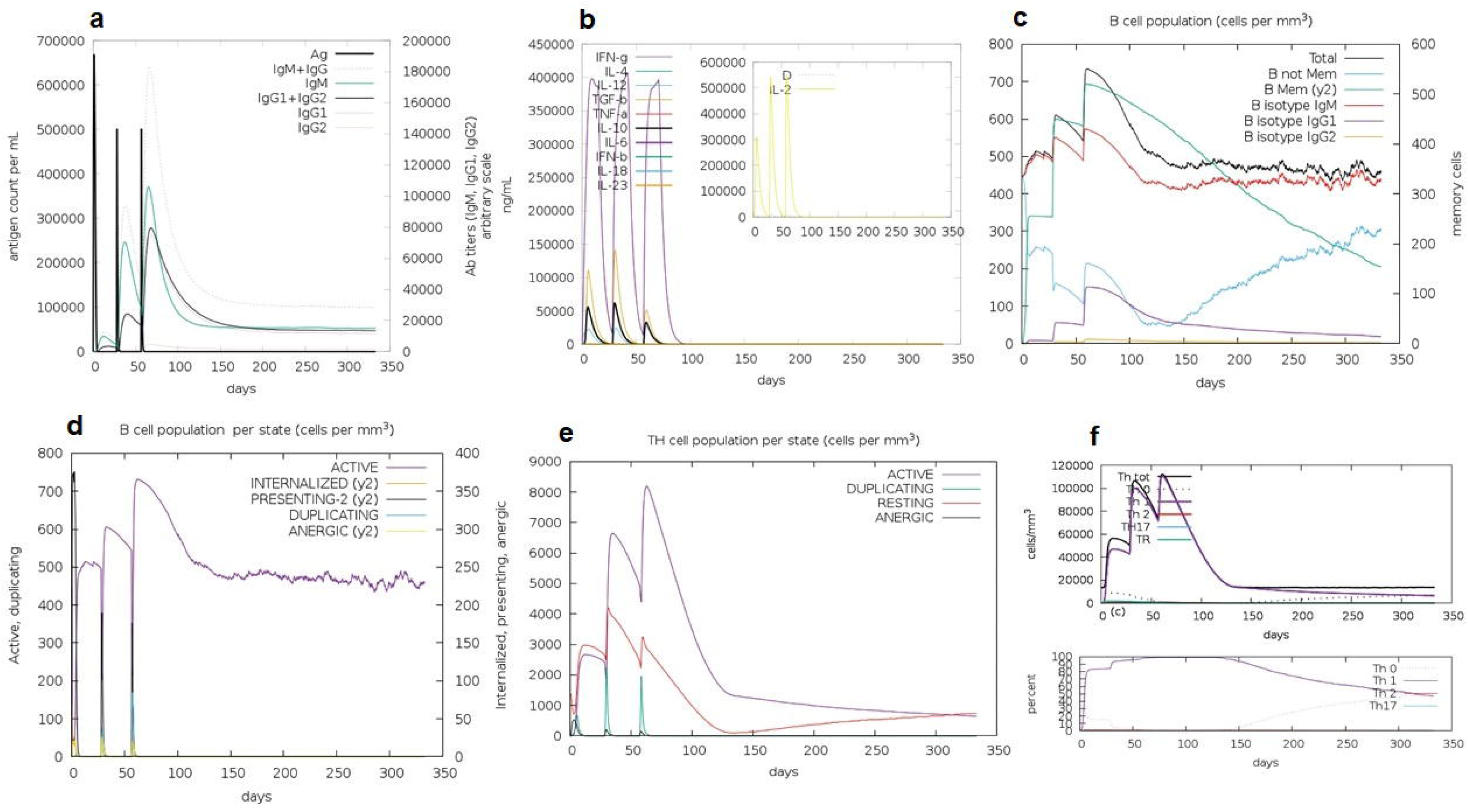
C-ImmSim presentation of an *in silico* immune simulation with the chimeric peptide. (a) The immunoglobulins and the immunocomplex response to antigen (CoV-RMEN) inoculations (black vertical lines); specific subclasses are indicated as colored peaks, (b) concentration of cytokines and interleukins, and inset plot shows danger signal together with leukocyte growth factor IL-2, (c) B-cell populations after three injections, (d) evolution of B cell, (e) T-helper cell populations per state after injections, and (f) evolution of T-helper cell classes with the course of vaccination.

### Molecular docking of CoV-RMEN with immune receptors (TLR3 and TLR4)

The ClusPro server was used to determine the protein binding and hydrophobic interaction sites on the protein surface. The immune responses of TLR3 and TLR4 against vaccine construct (CoV-RMEN) were estimated by analyzing the overall conformational stability of vaccine protein-TLRs docked complexes. The active interface aa residues of refined complexes of CoV-RMEN and TLRs were predicted (Fig. 8, Table 3). The relative binding free energies (ΔG) of the protein-TLRs complexes were significantly negative (Table 3) which suggest that the interaction of the chimeric protein might favor stimulation of the TLR receptors. Interface contacts (IC) per property (ICs charged-charged: 16, ICs charged-polar: 22, ICs charged-apolar: 26, polar-polar: 6, ICs polar-apolar: 25 and apolar-apolar: 29) were for the vaccine protein-TLR3 complex. Also, vaccine protein-TLR4 complex showed similar (ICs) per property (ICs charged-charged: 5, ICs charged-polar: 11, ICs charged-apolar: 30, polar-polar: 4, ICs polar-apolar: 31 and apolar-apolar: 39). These data validate the selected docked complexes that may result high binding among TLRs and the chimeric protein (Fig. 8).

**Fig. 8.**
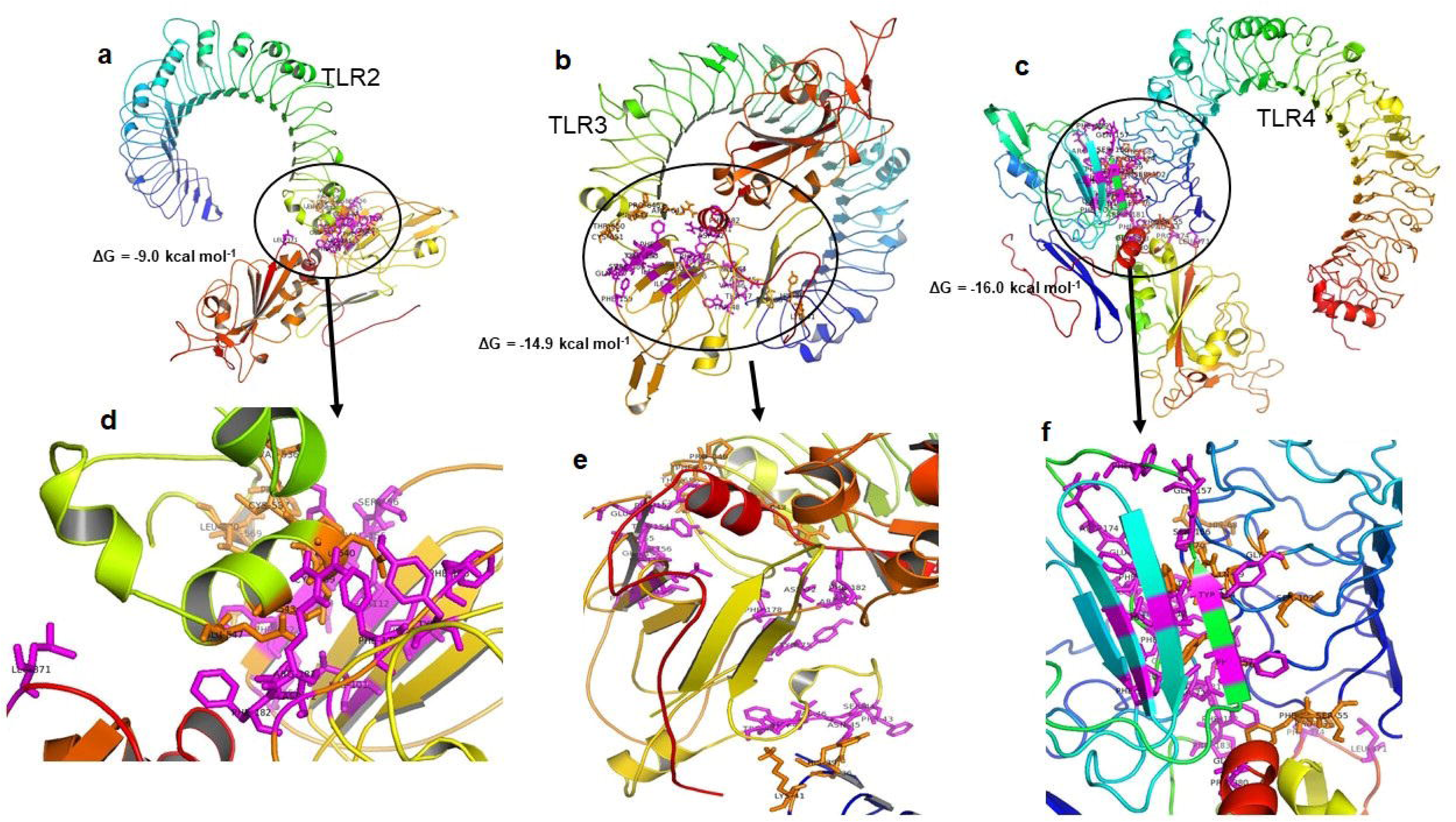
Molecular docking of CoV-RMEN vaccine with immune receptors (TLR2, TLR3 and TLR4). Docked complexes for (a) CoV-RMEN and TLR2, (b) CoV-RMEN and TLR3, and (c) CoV-RMEN and TLR4. Magnified interfaces of the complexes are figured to (d), (e) and (f) respectively. Active residues of CoV-RMEN colored magenta, and of TLRs colored orange with stick view. ΔG represents the binding affinity of the complexes.

**Table 3:** Active interface amino acid residues and binding scores among Toll Like Receptors (TLRs) and the constructed vaccine CoV-RMEN.

| Active residues of TLRs | | Active residues of CoV-RMEN | HADDOCK score | $\Delta G$ (kcal mol<sup>-1</sup>) |
| --- | --- | --- | --- | --- |
| **TLR-2** | V536, C537, S538, C539, E540, S543, E547, P567, R569, L570 | D72, Y75, L101, I103, I112, C150, F152, E153, Y154, V155, S156, F176, F178, R181, F182, L371 | -30.4 | -9.0 |
| **TLR-3** | D36, H39, K41, R643, F644, P646, F647, T650, C651, E652, S653, I654, W656, F657, V658, N659, W660, I661, N662, E663 | F43, S44, N45, V46, T47, W48, D72, Y75, F76, L101, I103, I112, F152, E153, Y154, V155, S156, Q157, F159, F178, R181, F182, L371 | -47.2 | -14.9 |
| **TLR-4** | P53, F54, S55, H68, G70, Y72, S73, F75, S76, Q99, S102, G124, | G39, L40, D72, Y75, F76, F90, L101, I103, I112, F152, Y154, S156, Q157, F159, R174, E175, F176, F178, R181, F182, P183, L371, P374, P380, G381 | -52.1 | -16.0 |

### Codon optimization of the CoV-RMEN

The high yield expression of a recombinant protein is always remained as a challenge, whereas heterologous prokaryotic expression vector can potentiate a robust, low cost and user-friendly technology for large scale vaccine production. For this reason, the recombinant protein encoding sequence need to be optimized for escalation of protein expression and downstream processing for a final product development. In order to optimize codon usage of the vaccine construct CoV-RMEN in *E. coli* (strain K12) for maximal protein expression, Rare Codon Analysis Tool (GenScript) was used. The length of the optimized codon sequence was 1,251 nucleotides. The Codon Adaptation Index (CAI) of the optimized nucleotide sequence was 0.87, and the average GC content of the adapted sequence was 50.26% showing the possibility of good expression of the vaccine candidate in the *E. coli* host. The optimal percentage range of GC content is between 30% and 70% (Fig. 9 a,b,c).

**Fig. 9.**
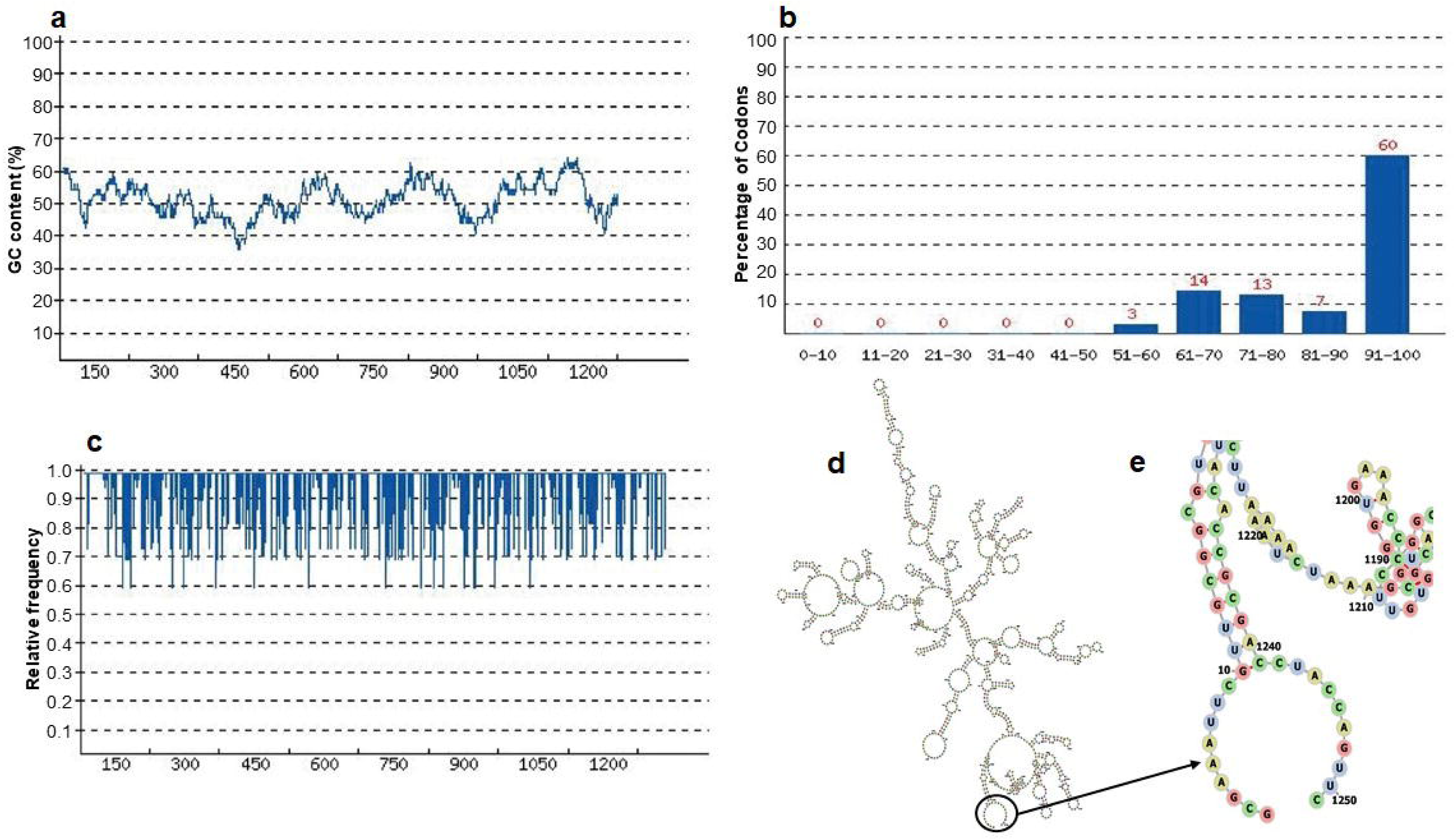
Codon optimization and mRNA structure of CoV-RMEN gene for expression in *E. coli*. (a) GC curve (average GC content: 50.26%) of the optimized CoV-RMEN gene, (b) percentage distribution of codons in computed codon quality groups, (c) relative distribution of codon usage frequency along the gene sequence to be expressed in *E. coli*, and codon adaptation index (CAI) was found to be 0.87 for the desired gene, (d) secondary structure and stability of corresponding mRNA, and (e) resolved view of the start region in the mRNA structure of CoV-RMEN.

### Prediction of mRNA secondary structure of the CoV-RMEN

The evaluation of minimum free energy for 25 structures of chimeric mRNA, the optimized sequences carried out by the ‘Mfold’server. The results showed that ΔG of the best predicted structure for the optimized construct was ΔG = −386.50 kcal/mol. The first nucleotides at 5’ did not have a long stable hairpin or pseudoknot. Therefore, the binding of ribosomes to the translation initiation site, and the following translation process can be readily accomplished in the target host. These outcomes were in the agreement with data obtained from the ‘RNAfold’web server (Fig. 9 d,e) where the free energy was −391.37 kcal/mol.

### Expression of CoV-RMEN with SUMO-fusion

After codon optimization and mRNA secondary structure analysis, the sequence of the recombinant plasmid was designed by inserting the adapted codon sequences into pETite vector (Lucigen, USA) using SnapGene software (Fig. 10). The utilization of pETite vector containing SUMO (Small Ubiquitin-like Modifier) tag and 6x-His tag will facilitate both the solubilization and affinity purification of the recombinant protein^29^. After successful cloning of the gene, the recombinant plasmid can be propagated efficiently using *E. coli* cells, and subsequent protein expression can be performed in *E. coli* K-12 strain using IPTG (Isopropyl β-d-1-thiogalactopyranoside) induction and cultivation at 28 °C as also reported earlier^29^.

**Fig. 10.**
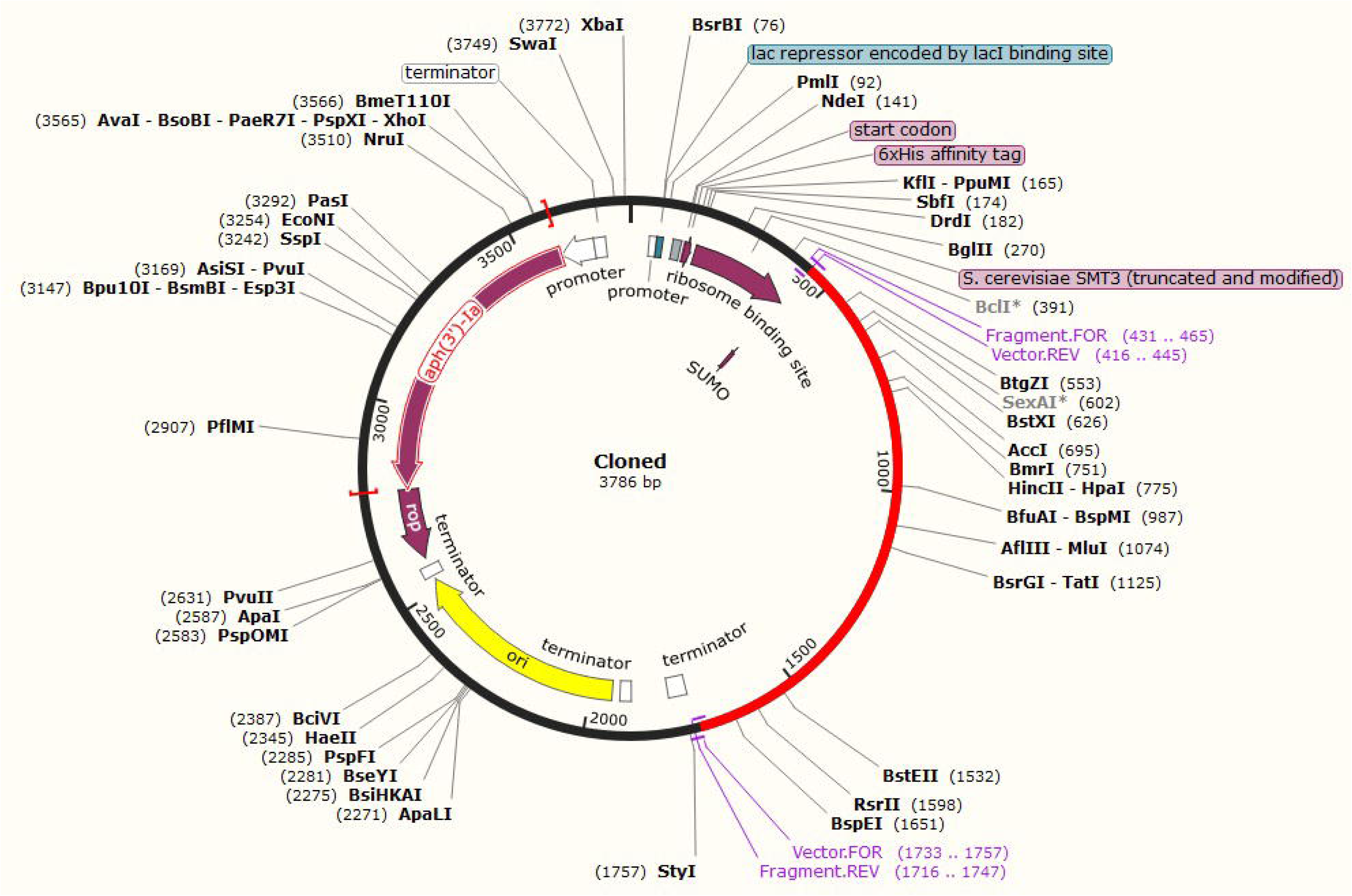
*In silico* fusion cloning of the CoV-RMEN. The final vaccine candidate sequence was inserted into the pETite expression vector where the red part represents the gene coding for the predicted vaccine, and the black circle represents the vector backbone. The six His-tag and SUMU-tag are located at the Carboxy-terminal end.

## Discussion

The causative agent of the ongoing COVID-19 pandemic caused by SARS-CoV-2, the seventh coronavirus with high zoonotic importance and transmission rate, has proved to be transmitted through direct and indirect contact along with airborne transmission which contribute very potentially for community transmission^30–32^. Since the emergence of SARS-CoV-2 in 2019 in Wuhan, China^31, 32^, it has exceeded both SARS-CoV and MERS-CoV in its rate of transmission among humans^30^, and therefore, effective measures counteracting its infection have become a major research focus. Scientific efforts have been going on extensively to learn about the virus, its biological properties, epidemiology, and pathophysiology^7^. At this early stage, different vaccine development efforts have been going on globally along with the novel drug development strategies^33^.

Development of an effective live attenuated vaccine for viral pathogens has been the most practiced strategy for prevention. But due to its extremely high contagious nature and ability for community transmission, this novel SARS-CoV-2 attenuated vaccine research essentially requires high biosafety level for researchers and work stations, which may eventually make it difficult to avail the vaccine for mass population of all the countries. Although live attenuated vaccine is often very immunogenic, however, the risk of reversion to a virulent virus has limited its usage as SARS-CoV and MERS-CoV vaccine^22^. Unlike other types of vaccines, chimeric peptide vaccine production does not involve virus replication, therefore reduce the cost of production. Hence, a low cost strategy should be adopted for developing a highly demanded vaccine for the mankind. Heterologous expression of any vaccine candidate protein has very promising scopes for developing such low cost vaccine, providing that all essential properties for antigenicity, immunogenicity and functional configuration are being conserved to mimic the structural and functional property of the actual antigen^34^. Construction of a vaccine candidate with multiple potential epitopes can obviously potentiate the multi-valency of the antigen to develop immune response against a number of epitopes of any pathogen. Also, rational engineering of epitopes for increased potency and magnitude, ability to enhance immune response in conserved epitopes, increased safety and absence of unnecessary viral materials and cost effectiveness all these cumulatively include potential benefit to multi-epitope recombinant protein based vaccine^20^. This study was designed to assist with the initial phase of multi-epitope vaccine candidate selection. Thereby, safe and effective vaccine development by providing recommendations of epitopes that may potentially be considered for incorporation in vaccine design for SARS-CoV-2.

Pathophysiology and virulence mechanisms of coronaviruses (CoVs), and therefore similarly of SARS-CoV-2 have links to the function of the structural proteins such as spike (S) glycoprotein, envelop (E), and membrane (M) proteins^34, 35^. Across the CoV families, the antibodies against the S glycoprotein, M and E proteins of SARS-CoV-2 would provide immunity to the infection^2, 12, 22^. Therefore, this work focused on the *in silico* design and development of a potential multi-epitope vaccine for SARS-CoV-2 selecting epitopes and domains from the S, E and M proteins.

Among the structural elements of CoVs, the spike S glycoproteins are composed of two major domains, the RBD and NTD. Furthermore, the homotrimeric S proteins of the SARS-CoV-2, SARS-CoV^9, 10^ and MERS-CoV^5^ comprises three chains: the chain A, B and C on the viral surface, guiding the link to host receptors. Unlike the full-length S protein of the CoVs, the RBD and NTD segments possessed the critical neutralizing domains but lacking the non-neutralizing immunodominant region^6, 10, 11, 22^. Therefore, considering the safety and effectiveness perspectives, the RBD and NTD are more promising candidates in the development of SARS-CoV-2 vaccines over the full-length S protein. The presence of E and M proteins on the envelope can augment the immune response against SARS-CoV^2, 36^ and thus, considered for suitable candidate for vaccine development^7, 12, 22^. Therefore, antibodies against the S, M and E proteins of SARS-CoV-2 would provide protective immunity to the infection^6, 7, 11, 12, 22^.

Immuno-informatics analyses of the selected RBD and NTD regions indicated that these contain ample amount of high-affinity B-cell, MHC Class I, MHC Class II and interferon-γ (IFN-γ) epitopes with high antigenicity scores. Moreover, membrane B-cell epitope (MBE) and envelop B-cell epitope (EBE) were selected from the M and E proteins, respectively for the final vaccine construct to enhance the overall stability, immunogenicity and antigenicity of the chimeric vaccine candidate, the CoV-RMEN^21^. The HLA alleles maintain the response to T-cell epitopes, and in different ethnic communities and geographical regions, these alleles are highly polymorphic. T-cell epitopes from RBD and NTD regions having high interaction with more HLA alleles were found to cover more than 98% of the world population with different ethnic groups^6, 37, 38^. Moreover, the molecular docking between T-cell epitopes and their respective HLA alleles revealed their highly significant binding affinity reflecting the immune activation of B- and T-cells^23^.

Vaccine design is improved through the use of specialized spacer sequences^39^. To designing the CoV-RMEN (vaccine candidate) GG and EGGE linkers were incorporated between the predicted epitopes to produce sequences with minimized junctional immunogenicity, thereby, allowing the rational design construction of a potent multi-epitope vaccine^21, 38^. The molecular weight of our vaccine candidate, the CoV-RMEN is 46.8 kDa with a predicted theoretical pI of 8.71, indicating that the protein is basic in nature. Also, the predicted instability index indicates that the protein will be stable upon expression, thus further strengthening its potential for use. The aliphatic index showed that the protein contains aliphatic side chains, indicating potential hydrophobicity. All these parameters indicate that the recombinant protein is thermally stable, hence would be best suited for use in different endemic areas worldwide^6, 21^.

The knowledge of secondary and tertiary structures of the target protein is essential in vaccine design^39, 40^. Secondary structure analysis of the CoV-RMEN indicated that the protein consisted of 43.2% alpha helix, 67.4% beta sheet, and 12% turns with only 2 residues disordered. Natively unfolded protein regions and alpha-helical and beta sheet peptides have been reported to be important forms of structural antigens^21, 40^. These two structural forms (secondary and tertiary), when tested as the synthetic peptides, have the ability to fold into their native structure,hence, could be recognized by naturally induced antibodies in response to infection^2^. The tertiary (3D) structure of the vaccine candidate improved markedly after the refinement, and showed desirable properties based on Ramachandran plot predictions. The Ramachandran plot shows that most of the residues are found in the favoured and allowed regions (94.7%) with very few residues in the outlier region; this indicates that the quality of the overall model is satisfactory^21, 23^. The lack of allergenic properties of the CoV-RMEN further strengthens its potential as a vaccine candidate. Strikingly, the multi-epitope peptide we designed and constructed (CoV-RMEN), showed higher antigenicity scores on Vaxijen v2.0 (0.450 with a virus model at a threshold of 0.4) and ANTIGENpro (0.875) gave an antigenic score almost twice that of Ov-103 on ANTIGENpro (0.93 for the chimeric peptide and 0.59 for Ov-103) supporting that this multiple-epitopes carrying vaccine candidate would be poorly immunogenic and require coupling to adjuvants. However, the designed protein must show similar antigenicity scores with and without the addition of an adjuvant sequence^21^.

It has been reported that immunity to viral infections is dependent on both B- and T- cells^20, 21^. Toll-Like Receptors (TLRs) recognize Pathogen Associated Molecular Patterns (PAMPs) from viral components or replication intermediates, resulting in signaling cascades that initiate an antiviral state in cells as a result of infection^41^. However, it is likely that TLRs might play an important role in the innate immune response to SARS-CoV-2 infection^42^. Though, we did not use any adjuvants, it is reported that TLRs (TLR3 and TLR4) can effectively bind with spike protein of the CoV^43^. The molecular docking analysis of the CoV-RMEN through HADDOCK showed that the chimeric protein can establish stable protein-protein interactions with TLRs (TLR-3,TLR-4)^41^ indicating efficient activation of these surface molecules which is very crucial for immune activation of dendritic cells for subsequent antigen processing and presentation to CD4+ and CD8+ T-cells via MHC-II and MHC-1, respectively corroborating the findings of different earlier studies^11, 20, 21^. TLR-4 might bind to S protein leading to the activation of the MyD88 dependent signaling pathway, which ultimately releases proinflammatory cytokines^35^. Immune simulation of the designed CoV-RMEN showed results consistent with typical immune responses, and there was a general increase in the generated immune responses after repeated antigen exposures. The comprehensive immune response developed after repeated antigen exposure reflected a perfect host defence development with the uplifted primary immune response and subsequent secondary immune response resulting in immune memory development^2^. In the present study, the development of memory B-cells and T-cells was evident, with memory in B-cells lasting for several months. Helper T-cells were particularly stimulated. The engrossing findings of the study is the development of Th1 response which enhance the growth and proliferation of B-cells augmenting the adaptive immunity^44^. The antiviral cytokine IFN-γ and cell stimulatory IL-2 level significantly increased which also contribute to the subsequent immune response after vaccination in host^2^. This indicates high levels of helper T-cells and consequently efficient antibody production, supporting a humoral response^21, 45^. The Simpson index, D for investigation of clonal specificity suggests a possible diverse immune response. This is plausible considering the generated chimeric peptide is composed of sufficient B- and T-cell epitopes.

One of the first steps in validating a candidate vaccine is to screen for immunoreactivity through serological analysis. This requires the expression of the recombinant protein in a suitable host. *Escherichia coli* expression systems are the preferred choice for the production of recombinant proteins^33^. In order to achieve high-level expression of our recombinant vaccine protein in *E. coli* (strain K12), codon optimization was carried out. Stable mRNA structure, codon adaptability index (0.87), and the GC content (50.26%) were favourable for high-level expression of the protein in bacteria. The next step projected at the moment, is to express this peptide in a bacterial system and perform the various immunological assays needed to validate the results obtained here through immuno-informatics analyses.

## Conclusions

COVID-19 is reaching entire globe and emergence of the causative agent SARS-CoV-2 kept us nowhere as we are racing for treatment and prevention. This multi-epitope chimera that we named as CoV-RMEN possess domains of RBD and NTD segments of S, M and E proteins all of which showed significant antigenic properties compared to any other viral proteins. This chimera also includes potential CTL, HTL and B-cell epitopes to ensure humoral as well as cellular immune response and the optimal expression and stability of the chimera was validated. With multiple limitations and high cost requirements for the attenuated vaccine preparation for contagious agents like SARS-CoV-2, this chimeric peptide vaccine candidate gives us the hope to ensure it’s availability and relatively cheap option to reach entire world. This CoV-RMEN can be very effective measure against COVID-19 to reach globally. Hence, this could be cloned, expressed and tried for *in vivo* validations and animal trials at the laboratory level.

## Methods

### Sequence retrieval and structure generation

A total of 250 partial and complete genome sequences of SARS-CoV-2 were retrieved from NCBI (Supplementary Table 5). We aligned these sequences through MAFFT online server (https://mafft.cbrc.jp/alignment/server/) using default parameters, and Wu-Kabat protein variability was analyzed in Protein variability server (http://imed.med.ucm.es/PVS/) with respect to NCBI (National Center for Biotechnology Information, https://www.ncbi.nlm.nih.gov/protein) reference genome (Accession no : NC_045512.2). Despite having minor heterogeneity in spike glycoprotein (S), membrane (M) and envelop (E) proteins, the NCBI reference genome (Accession no: NC_045512.2) of SARS-CoV-2 from Wuhan, China was finally selected for domains and epitopes selection, secondary and tertiary (3D) structure prediction, antigenicity and allergenicity assessment, refinement and validation, molecular docking, cDNA and mRNA analysis for cloning and expression^20, 21^. Moreover, two reference genome sequences of SARS-CoV (NC_004718.3) and MERS-CoV (NC_019843.3) were also retrieved from the NCBI database to reveal the structural heterogeneity of S protein. Multiple sequence alignment of the S proteins of the viruses was performed by CLUSTALW^45^. Since the S glycoprotein of coronaviruses is important for viral attachment to host cell, antibody against this protein acts as inhibitor^30^, therefore, S proteins of SARS-CoV, MERS-CoV and SARS-CoV-2 were structurally compared using SWISS homology modeling^46^ based on the protein databank (PDB) templates 6acd, 5w9h and 6vsb, respectively. The models were then optimized by energy minimization using the GROMOS96 program^27^ implemented within the Swiss-PdbViewer, version 4.1.0^47^. The Ramachandran plots of the derived models were evaluated using a PROCHECK (version 3.5)-server to check the stereochemical properties of the modeled structures^48^. Finally, the homology models were structurally aligned using PyMOL^49^, and observed for heterogeneity in the conformations. The physiochemical properties of the S glycoprotein of the SARS-CoV-2, SARS-CoV and MERS-CoV was computed using the Protparam ExPasy-server^50^.

### Linear B-cell epitopes prediction

We employed both structure and sequence-based methods for B-cell epitopes prediction. Conformational B-cell epitopes on the S protein were predicted by Ellipro (Antibody Epitope Prediction tool; http://tools.iedb.org/ellipro/) available in IEDB analysis resource^51^ with the minimum score value set at 0.4 while the maximum distance selected as 6 Å. The ElliPro allows the prediction and visualization of B-cell epitopes in a given protein sequence or structure. The ElliPro method is based on the location of the residue in the protein’s three-dimensional (3D) structure. ElliPro implements three algorithms to approximate the protein shape as an ellipsoid, calculate the residue protrusion index (PI), and cluster neighboring residues based on their protrusion index (PI) value. The residues lying outside of the ellipsoid covering 90% of the inner core residues of the protein score highest PI of 0.9^23^. Antigenicity of full-length S (spike glycoprotein), M (membrane protein) and E (envelope protein) proteins was predicted using VaxiJen v2.0 (http://www.ddg-pharmfac.net/vaxijen/VaxiJen/VaxiJen.html)^52^. The linear B-cell epitopes of RBD and NTD regions of S protein, full length E and M proteins were predicted by “BepiPred Linear Epitope Prediction” (propensity scale method such as hidden Markov model)^53^ and ABCPred under default parameters^54^. To find out the most probable peptide-based epitopes with better confidence level, selected peptides were further tested using VaxiJen antigenicity scores^51^. The Kolaskar and Tongaonkar antigenicity profiling from IEDB analysis resource was also used for RBD and NTD segments, E and M proteins^55^.

### Screening for T-cell epitopes

Immune Epitope Database (IEDB) tool peptide binding to MHC class I molecules and Proteasomal cleavage/TAP transport/MHC class I combined predictor (http://tools.iedb.org/main/tcell/) were used to screen CTL epitopes, proteasomal cleavage and transporter associated with antigen processing (TAP) with all parameters set to default. Threshold for strong binding peptides (IC_50_) was set at 50 nM to determine the binding and interaction potentials of helper T-cell epitope peptide and major histocompatibility complex (MHC) class I allele^20, 23^. The total score generated by the tool is a combined score of proteasome, major histocompatibility complex (MHC), TAP (N-terminal interaction), processing analysis scores. The HTL epitopes were screened using the IEDB tool “MHC-II Binding Predictions” (http://tools.iedb.org/mhcii/). The tool generated the median percentile rank for each predicted epitope through combination of three methods (Combinatorial Library, SMM-align, and Sturniolo) by comparing the score of peptide against the scores of other random five million 15-mer peptides from the SwissProt database^28^. Top five HLA epitopes for each RBD and NTD segments were docked against the respective HLA (MHC-□ and MHC-□) allele binders by interaction similarity-based protein-peptide docking system GalaxyPepDock of the GalaxyWeb, docked HLA-epitope complexes were refined in GalaxyRefineComplex and binding affinity (ΔG) was determined PROtein binDIng enerGY prediction (PRODIGY) tool^56^.

### Population coverage by CTL and HTL epitopes

Due to the MHC restriction of T-cell response, the peptides with more different HLA binding specificities mean more population coverage in defined geographical regions and ethnic groups where-and-to-whom the peptide-based vaccine might be employed. The predicted T-cell epitopes were shortlisted based on the aligned Artificial Neural Network (ANN) with half-maximal inhibitory concentration (annIC_50_) values up to 50 nM. The IEDB “Population Coverage” tool (http://tools.iedb.org/population/)^57^ was used to determine the world human population coverage by the shortlisted CTL and HTL epitopes. We used multi-epitopes involving both CTL and HTL epitopes to have the higher probability of larger human population coverage worldwide. We used OmicsCircos to visualize the association between world population and different ethnic groups^58^.

### IFN-**γ**-inducing epitope prediction

Potential IFN-γ epitopes of all the selected antigenic sites of RBD, NTD, envelop protein B-cell epitope (EBE), and membrane protein B-cell epitope (MBE) were predicted by “IFNepitope” server (http://crdd.osdd.net/raghava/ ifnepitope/scan.php). To identify the set of epitopes associated with MHC alleles that would maximize the population coverage, we adopted the “Motif and SVM hybrid” (MERCI: Motif-EmeRging and with Classes-Identification, and SVM) approach. The tool generates overlapping sequences from the query antigens for IFN-γ-inducing epitope prediction. The estimated population coverage represents the percentage of individuals within the population that are likely to elicit an immune response to at least one T cell epitope from the set, and finally the coverage was observed by adding epitopes associated with any of the remaining MHC alleles^2^. The prediction is based on a dataset of IFN-γ-inducing and IFN-γ-noninducing MHC allele binders^59^.

### Design and construction of multi-epitope vaccine candidate

The candidate vaccine design and construction method follows previously developed FMD peptide vaccine development protocol in Professor M Anwar Hossain’s laboratory (unpublished data, and under patent application in Bangladesh Application No. P/BD/2018/000280, date: October 1, 2018; India Patent Number: 201924036919, Date: September 13, 2019). The multi-epitope protein was constructed by positioning the selected RBD, NTD, MBE and EBE amino acid sequences linked with short, rigid and flexible linkers GG. To develop highly immunogenic recombinant proteins, two universal T-cell epitopes were used, namely, a pan-human leukocyte antigen DR-binding peptide (PADRE)^60^, and an invasin immunostimulatory sequence taken from *Yersinia* (Invasin)^61^ were used to the N and C terminal of the vaccine construct respectively, linked with EGGE. Here, acidic aa, Glutamate (E) was added to balance the ratio of acidic and basic amino acids. We denoted the vaccine candidate as CoV-RMEN. VaxiJen 2.0^52^ and ANTIGENpro (http://scratch.proteomics.ics.uci.edu/) web-servers were used to predict the antigenicity of the CoV-RMEN while the AllerTOP 2.0 (http://www.ddg-pharmfac.net/AllerTOP) and AllergenFP (http://ddg-pharmfac.net/AllergenFP/) web-servers were used to predict vaccine allergenicity.

### Physiochemical properties and solubility prediction of the CoV-RMEN

Various physiochemical parameters of the CoV-RMEN were assessed using the online web-server ProtParam^50^. These parameters included aa residue composition, theoretical pI, instability index, *in vitro* and *in vivo* half-life, aliphatic index, molecular weight, and grand average of hydropathicity (GRAVY). The solubility of the multi-epitope vaccine peptide was evaluated using the Protein-Sol server (https://protein-sol.manchester.ac.uk/).

### Secondary and tertiary structure prediction

Chou and Fasman secondary structure prediction server (CFSSP: https://www.biogem.org/tool/chou-fasman/) and RaptorX Property (http://raptorx.uchicago.edu/StructurePropertyPred/predict/)^62^, web-servers were used for secondary structure predictions. The RaptorX web-server (http://raptorx.uchicago.edu/)^62^ was used to predict the three-dimensional (3D) structure and binding residues of the chosen protein. Homology modelling of the CoV-RMEN was carried out using the protein homology/analogy recognition engine (Phyre2) (http://www.sbg.bio.ic.ac.uk/~phyre2/html/page.cgi?id=index) web-server.

### Refinement and validation of the tertiary structure of the CoV-RMEN

The 3D model obtained for the CoV-RMEN was refined in a three-step process, initially energy minimization using the GROMOS96 program^27^ implemented within the Swiss-PdbViewer (version 4.1.0)^47^. After energy minimization, the model was refined using ModRefiner (https://zhanglab.ccmb.med.umich.edu/ModRefiner/) and then GalaxyRefine server (http://galaxy.seoklab.org/cgi-bin/submit.cgi?type=REFINE). The construction and refinement of the CoV-RMEN was further assessed through ModRefiner server for atomic-level energy minimization. This results in improvements in both global and local structures, with more accurate side chain positions, better hydrogen-bonding networks, and fewer atomic overlaps^63^. The GalaxyRefine server was further used to improving the best local structural quality of the CoV-RMEN according to the CASP10 assessment, and ProSA-web (https://prosa.services.came.sbg.ac.at/prosa.php) was used to calculate overall quality score for a specific input structure, and this is displayed in the context of all known protein structures. The ERRAT server (http://services.mbi.ucla.edu/ERRAT/) was also used to analyze non-bonded atom-atom interactions compared to reliable high-resolution crystallography structures. A Ramachandran plot was obtained through the RAMPAGE server (http://mordred.bioc.cam.ac.uk/~rapper/rampage.php). The server uses the PROCHECK principle to validate a protein structure by using a Ramachandran plot and separates plots for Glycine and Proline residues^64^.

### Immune simulation

To further characterize the immunogenicity and immune response profile of the CoV-RMEN, *in silico* immune simulations were conducted using the C-ImmSim server (http://150.146.2.1/C-IMMSIM/index.php)^28^. All simulation parameters were set at default with time steps set at 1, 84, and 170 (each time step is 8 hours and time step 1 is injection at time = 0). Therefore, three injections were given at four weeks apart. The Simpson index, D (a measure of diversity) was interpreted from the plot.

### Molecular docking of the CoV-RMEN with TLRs

The generation of an appropriate immune response is dependent on the interaction between an antigenic molecule and a specific immune receptor like Toll Like Receptors (TLRs)^33^. Molecular docking of the CoV-RMEN with the TLR3 (PDB ID: 1ZIW) and TLR4 (PDB ID: 4G8A) receptor was performed using the High Ambiguity Driven DOCKing (HADDOCK, version 2.4)^65^ web-server in order to evaluate the interaction between ligand and receptor and consequently the development of an immune response. In order to predict such interaction, data-driven docking of designed chimeric protein and TLR3 and TLR4 complexes were performed^33^. In this regard, CPORT (https://milou.science.uu.nl/services/CPORT/) was used to predict active interface residues of the CoV-RMEN and TLRs. The HADDOCK server was then employed to perform docking simulations of the CoV-RMEN-TLRs complexes due to perform short MD simulations of docked complexes^56^. Finally, the binding affinities of the top-ranking docking pose of chimeric protein-TLRs complexes was predicted using the PRODIGY (PROtein binDIng enerGY prediction) (https://nestor.science.uu.nl/prodigy/) web-server^56^.

### Analysis of cDNA and mRNA for cloning and expression

Reverse translation and codon optimization were performed using the GenScript Rare Codon Analysis Tool (https://www.genscript.com/tools/rare-codon-analysis) in order to express the CoV-RMEN in a selected expression vector. Codon optimization was performed in order to express the final vaccine construct in the *E. coli* (strain K12). The analysis output includes the codon adaptation index (CAI) and percentage GC content, which can be used to assess protein expression levels. CAI provides information on codon usage biases; the ideal CAI score is 1.0 but >0.8 is considered a good score^66^. The GC content of a sequence should range between 30–70%. GC content values outside this range suggest unfavorable effects on translational and transcriptional efficiencies^22^. Stability of the mRNA was measured using two different tools namely RNAfold (http://rna.tbi.univie.ac.at/cgi-bin/RNAWebSuite/RNAfold.cgi) and the mfold (http://unafold.rna.albany.edu/?q=mfold) web-servers. To clone the optimized gene sequence of the final vaccine construct in *E. coli* into pETite vector (Lucigen, USA) through enzyme-free method. The primers for amplification of the DNA fragment must contain 18-nucletide short DNA sequence (5’-CGCGAACA-GATTGGAGGT-3’ in upstream of forward primer and 5’-GTGGCGGCCGCTCTATTA-3’ in upstream of reverse primer) to facilitate the enzyme-free cloning of the gene at the insertion site. Finally, the sequence of the recombinant plasmid was designed by inserting the adapted codon sequences into pETite vector using SnapGene software (from Insightful Science; available at snapgene.com).

## Author contributions

MSR, MNH, and MRI done the overall study and also draft the manuscript. SA drafted some parts of results and discussion. MNH finally compiled the manuscript. OS, and MAS helped in final discussion and reference indexing. ASMRUA, MMR, MS and MAH contributed intellectually to the interpretation and presentation of the results. MS and MAH developed the concept of peptide vaccine development. Finally, all authors have approved the manuscript for submission.

## Acknowledgements

The authors thank Joynob Akter Puspo, Masuda Akter and Israt Islam (MS student), and DR. Kazi Alamgir Hossain (PhD Fellow) of the Microbial Genetics and Bioinformatics Laboratory, Department of Microbiology, University of Dhaka for their cooperation, suggestions and overall encouragement during the preparation of the manuscript.

## Supplementary Material

Supplementary information supporting the findings of the study are available in this article as Supplementary Data files, or from the corresponding author on request.

---

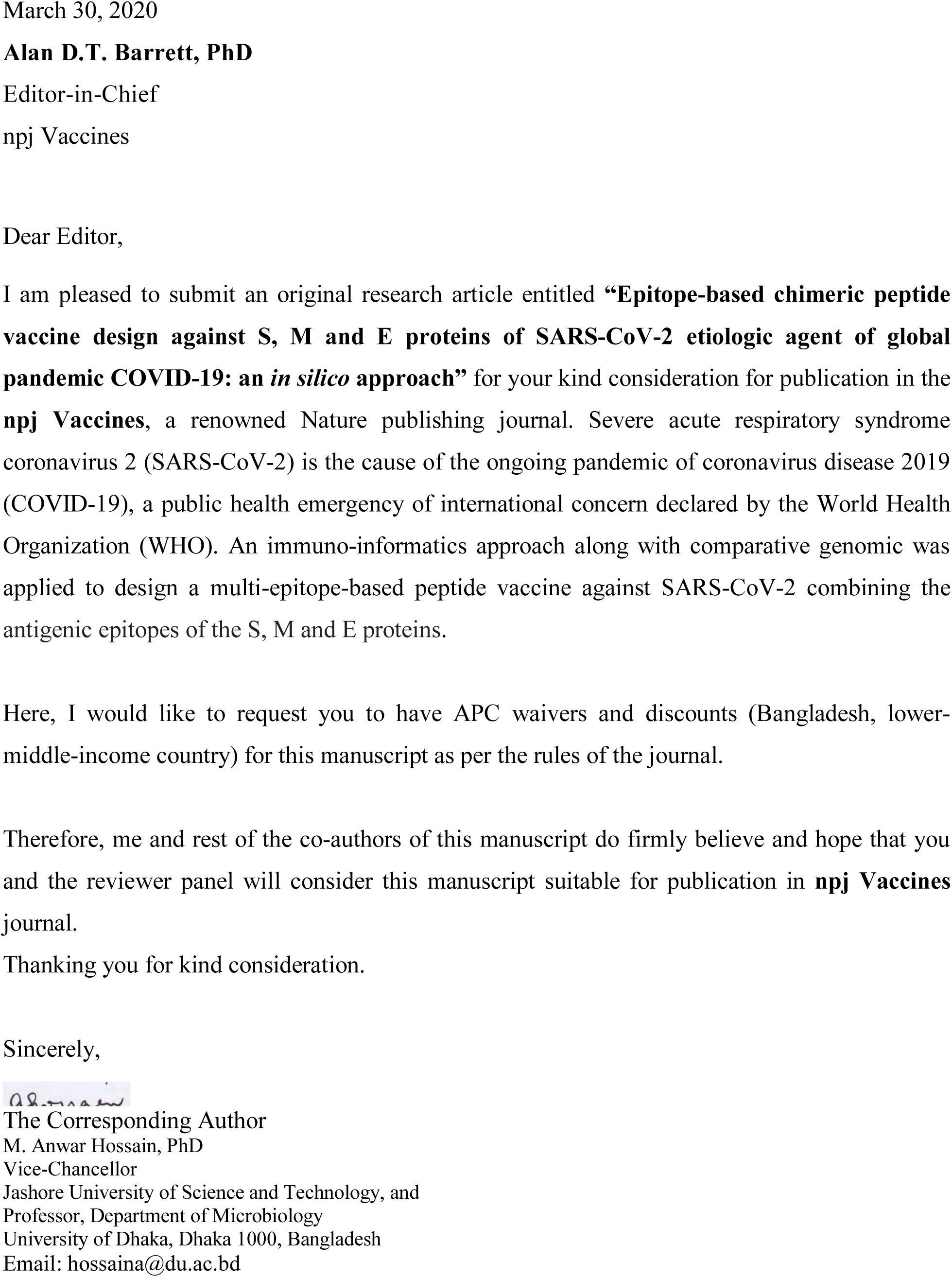

## References

1. WHO, 2020. Novel Coronavirus (2019-nCoV) situation reports - World Health Organization (WHO)

2. Almofti, Y. A., Abd-elrahman, K. A., Gassmallah, S. A. E. & Salih, M. A. Multi Epitopes Vaccine Prediction against Severe Acute Respiratory Syndrome (SARS) Coronavirus Using Immunoinformatics Approaches. *American J*. Microbiol. Res. 6(3), 94–114 (2018).

3. Wu, F. et al. “A new coronavirus associated with human respiratory disease in China.” Nat. 1–5 (2020)

4. Badawi, M. M. et al. *In Silico* Prediction of a Novel Universal Multi-epitope Peptide Vaccine in the Whole Spike Glycoprotein of MERS CoV. Am. J. Microbiol Res. 4(4), 101–21 (2016).

5. Pallesen, J. et al. Immunogenicity and structures of a rationally designed prefusion MERS-CoV spike antigen. Proc. Nat. Aca. Sci. 114(35), E7348–E7357 (2017).

6. ul Qamar, M. T., Saleem, S., Ashfaq, U. A., Bari, A., Anwar, F. & Alqahtani, S. Epitope based peptide vaccine design and target site depiction against Middle East Respiratory Syndrome Coronavirus: an immune-informatics study. J. Translational Med. 17(1), 362 (2019).

7. Ahmed, S. F., Quadeer, A. A. & McKay, M. R. Preliminary identification of potential vaccine targets for the COVID-19 coronavirus (SARS-CoV-2) based on SARS-CoV immunological studies. Viruses. 12(3), 254 (2020).

8. Abdelmageed, M. I. et al. Design of multi epitope-based peptide vaccine against E protein of human 2019-nCoV: An immunoinformatics approach. BioRxiv. (2020).

9. Song, W., Gui, M., Wang, X. & Xiang, Y. Cryo-EM structure of the SARS coronavirus spike glycoprotein in complex with its host cell receptor ACE2. PLoS Pathog. 14(8), e1007236 (2018).

10. Wrapp, D. et al. Cryo-EM structure of the 2019-nCoV spike in the prefusion conformation. Science 2020.

11. Shang, W. et al. The outbreak of SARS-CoV-2 pneumonia calls for viral vaccines. npj Vaccines 5, 18 (2020).

12. Gralinski, L. E. & Menachery, V. D. Return of the coronavirus: 2019-nCoV. Viruses 12, 135 (2020).

13. Zhou, H. et al. Structural definition of a neutralization epitope on the N-terminal domain of MERS-CoV spike glycoprotein. Nat. Commun. 10, 3068 (2019).

14. Rabi, F. A., Al Zoubi, M. S., Kasasbeh, G. A., Salameh, D. M. & Al-Nasser, A. D. SARS-CoV-2 and Coronavirus Disease 2019: What We Know So Far. Pathogens 9(3), 231 (2020).

15. Wang, N. et al. Structural Definition of a Neutralization-sensitive Epitope on the MERS-CoV S1-NTD. Cell Rep. 28(13), 3395–405 (2019).

16. Shi, S. Q. et al. The expression of membrane protein augments the specific responses induced by SARS-CoV nucleocapsid DNA immunization. Mol. Immunol. 43, 1791–1798 (2006).

17. Schoeman, D. & Fielding, B. C. Coronavirus envelope protein: current knowledge. Virol J. 16, 69 (2019).

18. Chan, J. F. et al. Genomic characterization of the 2019 novel human-pathogenic coronavirus isolated from a patient with atypical pneumonia after visiting Wuhan. Emerg. Microbes Infect. 9, 221–236 (2020).

19. Li, G., et al. Coronavirus infections and immune responses. J. Med. Virology 92(4), 424–432 (2020).

20. Shi, J. et al. Epitope-Based Vaccine Target Screening against Highly Pathogenic MERS-CoV: An *In Silico* Approach Applied to Emerging Infectious Diseases. PLoS One 10(12), e0144475 (2015).

21. Shey, R. A. et al. *In-silico* design of a multi-epitope vaccine candidate against onchocerciasis and related filarial diseases. Sci Rep. 9(1), 1–18 (2019).

22. Yong, et al. Recent Advances in the Vaccine Development Against Middle East Respiratory Syndrome-Coronavirus. Front. Microbiol. 10, 1781 (2019).

23. Srivastava, S. et al. Design of novel multi-epitope vaccines against severe acute respiratory syndrome validated through multistage molecular interaction and dynamics. J. Biomol. Struc. Dyn. 37(16), 4345–4360 (2019).

24. Sakib, M. S., Islam, M., Hasan, A. K. M. & Nabi, A. H. M. Prediction of epitope-based peptides for the utility of vaccine development from fusion and glycoprotein of nipah virus using in silico approach. Adv. Bioinform. (2014).

25. Adhikari, U. K., Tayebi, M. & Rahman, M. M. Immunoinformatics approach for epitope-based peptide vaccine design and active site prediction against polyprotein of emerging oropouche virus. J. Immunol. Res. (2018).

26. Hebditch, M., M. A. Carballo-Amador, S. Charonis, R. Curtis & J. Warwicker. Protein– Sol: a web tool for predicting protein solubility from sequence. Bioinform. 33(19), 3098–3100 (2017).

27. Van Gunsteren, W., S., et al. The GROMOS96 manual and user guide. (1996).

28. Rapin, N., Lund, O, Bernaschi, M & Castiglione, M. Computational immunology meets bioinformatics: the use of prediction tools for molecular binding in the simulation of the immune system. PLoS One 5(4) (2010).

29. Biswal, J. K., Bisht, P., Mohapatra, J. K., Ranjan, R., Sanyal, A. & Pattnaik, B. “Application of a recombinant capsid polyprotein (P1) expressed in a prokaryotic system to detect antibodies against foot-and-mouth disease virus serotype O.” J. Virol. Methods 215, 45–51 (2015).

30. Tai, W. et al. Characterization of the receptor-binding domain (RBD) of 2019 novel coronavirus: implication for development of RBD protein as a viral attachment inhibitor and vaccine. Cell. Mol. Immunol. (2020).

31. Zhou, P. et al. Discovery of a novel coronavirus associated with the recent pneumonia outbreak in humans and its potential bat origin. bioRxiv. (2020).

32. Hui, D. S., et al. The continuing 2019-nCoV epidemic threat of novel coronaviruses to global health—The latest 2019 novel coronavirus outbreak in Wuhan, China. Int. J. Inft. Dis. 91, 264–266 (2020).

33. Pei, H. et al. Expression of SARS-coronavirus nucleocapsid protein in *Escherichia coli* and *Lactococcus lactis* for serodiagnosis and mucosal vaccination. Applied Microbiol. Biotech. 68(2), 220–227 (2005).

34. Pang, J. et al. Potential rapid diagnostics, vaccine and therapeutics for 2019 novel Coronavirus (2019-ncoV): a systematic review. J. Clinical Med. 9(3), 623 (2020).

35. Frieman, M. et al. “SARS coronavirus and innate immunity.” Virus Research. 133(*1*), 101–112 (2018).

36. Millet, J. K. & Whittaker, G. R. “Host cell proteases: Critical determinants of coronavirus tropism and pathogenesis.” Virus Res. 202, 120–134 (2015).

37. Jaimes, J. A., Andre, N. M., Millet, J. K. & Whittaker, G. R. Structural modeling of 2019-novel coronavirus (nCoV) spike protein reveals a proteolytically-sensitive activation loop as a distinguishing feature compared to SARS-CoV and related SARS-like coronaviruses. arXiv. (2020).

38. Huang, H. et al. CD4(+) Th1 cells promote CD8(+) Tc1 cell survival, memory response, tumor localization and therapy by targeted delivery of interleukin 2 via acquired pMHC I complexes. Immunology 120, 148–159 (2007).

39. Mohammad, N. et al. *In silico* design of a chimeric protein containing antigenic fragments of Helicobacter pylori; a bioinformatic approach. The Open Microbiol. J. 10, 97 (2016).

40. Meza, B., Ascencio, F., Sierra-Beltrán, A. P., Torres, J. & Angulo, C. A novel design of a multi-antigenic, multistage and multi-epitope vaccine against *Helicobacter pylori*: an *in silico* approach. Infect. Genet. Evol. 49, 309–317 (2017).

41. Totura, A. L. et al. Toll-like receptor 3 signaling via TRIF contributes to a protective innate immune response to severe acute respiratory syndrome coronavirus infection. MBio 6(3), e00638–15 (2015).

42. Mosaddeghi, P., et al. Therapeutic approaches for COVID-19 based on the dynamics of interferon-mediated immune responses. (2020).

43. Zander, R. A. et al. Th1-like plasmodium-specific memory CD4+ T cells support humoral immunity. Cell Rep. 21(7), 1839–1852 (2017).

44. Carvalho, L. H. et al. IL-4-secreting CD4+ T cells are crucial to the development of CD8+ T-cell responses against malaria liver stages. Nat. Med. 8, 166–70 (2002).

45. Hoque, M. N. et al. Metagenomic deep sequencing reveals association of microbiome signature with functional biases in bovine mastitis. Sci. Rep. 9, 13536 (2019).

46. Waterhouse, A. et al. “SWISS-MODEL: homology modelling of protein structures and complexes.” Nucleic Acids Res. 46(W1), W296–W303 (2018).

47. Guex, N. & Peitsch, M. C. “SWISS-MODEL and the Swiss-Pdb Viewer: an environment for comparative protein modeling.” Electrophoresis 18(15), 2714–2723 (1997).

48. Laskowski, R. A., MacArthur, M. W., Moss, D. S. & Thornton, J. M. “PROCHECK: a program to check the stereochemical quality of protein structures.” J. Applied Crystal 26(2), 283–291 (1993).

49. Faure, G. et al. iPBAvizu: a PyMOL plugin for an efficient 3D protein structure superimposition approach. Source Code Biol. Med. 14, 5 (2019).

50. Gasteiger, E. et al. Protein identification and analysis tools on the ExPASy server. *The Proteomics Protocols Handbook*, Springer: 571–607 (2005).

51. Kringelum, J. V., Lundegaard, C., Lund, O. & Nielsen, M. Reliable B cell epitope predictions: Impacts of method development and improved benchmarking. PLoS Comput. Biol. 8(12), e1002829 (2012).

52. Doytchinova, I. A. & Flower, D. R. VaxiJen: a server for prediction of protective antigens, tumour antigens and subunit vaccines. BMC Bioinformatics 8(1), 4 (2007).

53. Larsen, J. E. P., Lund, O. & Nielsen, M. Improved method for predicting linear B-cell epitopes. Immunome Res. 2(1), 2 (2006).

54. Saha, S. & Raghava G. P. S. “Prediction of continuous BLJ using recurrent neural network.” Proteins: Structure, Function, Bioinformatics 65(1), 40–48 (2006).

55. Kolaskar, A. & Tongaonkar, P. C. “A semi empirical method for prediction of antigenic determinants on protein antigens.” FEBS Letters 276(1-2), 172–174 (1990).

56. Xue, L. C., Rodrigues, J. P., Kastritis, P. L., Bonvin, A. M. & Vangone, A. “PRODIGY: a web server for predicting the binding affinity of protein–protein complexes.” Bioinformatics 32(23), 3676–3678 (2016).

57. Bui, H. H., Sidney, J., Dinh, K., Southwood, S., Newman, M. J & Sette, A. Predicting population coverage of T-cell epitope-based diagnostics and vaccines. BMC Bioinformatics 7(1),153 (2006).

58. Hoque, M. N. et al. Resistome diversity in bovine clinical mastitis microbiome, a signature concurrence. bioRxiv, 829283 (2019).

59. Dhanda, S. K., Vir, P. & Raghava, G. P. “Designing of interferon-gamma inducing MHC class-II binders.” Biology Direct 8(1), 30 (2013).

60. Agadjanyan, M. G. et al. “Prototype Alzheimer’s disease vaccine using the β-amyloid and promiscuous T cell epitope pan HLA DR-binding peptide.” The J. Immunology 174(3), 1580–1586 (2005).

61. Li, H. et al. Co-expression of the C-terminal domain of *Yersinia enterocolitica* invasin enhances the efficacy of classical swine-fever-vectored vaccine based on human adenovirus. J. Biosci. 40, 79–90 (2015).

62. Källberg, M., Margaryan, G., Wang, S., Ma, J. & Xu, J. RaptorX server: a resource for template-based protein structure modeling. In Protein Structure Prediction (pp. 17–27), Humana Press, New York, NY. 2014.

63. Xu, D. & Zhang, Y. “Improving the physical realism and structural accuracy of protein models by a two-step atomic-level energy minimization.” Biophysical J. 101(10), 2525–2534 (2011).

64. Lovell, S. C. et al. “Structure validation by Cα geometry □, ψ and Cβ deviation.” Proteins: Structure, Function, Bioinformatics 50(3), 437–450 (2003).

65. De Vries, S. J., Van Dijk, M. & Bonvin, A. M. The HADDOCK web server for data-driven biomolecular docking. Nat. Protoc. 5(5), 883 (2010).

66. Morla, S., Makhija, A. & Kumar, S. “Synonymous codon usage pattern in glycoprotein gene of rabies virus.” Gene 584(1), 1–6 (2016).

