## Supplementary figures and images for "Epitope-based chimeric peptide vaccine design against S, M and E proteins of SARS-CoV-2 etiologic agent of global pandemic COVID-19: an *in silico* approach"

### Supplementary Fig. 1

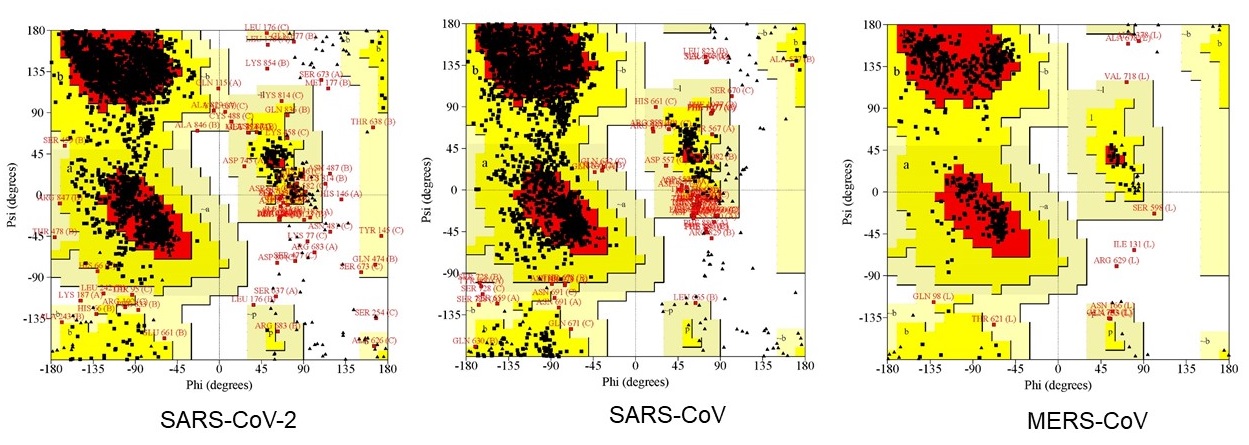

### Supplementary Fig. 2

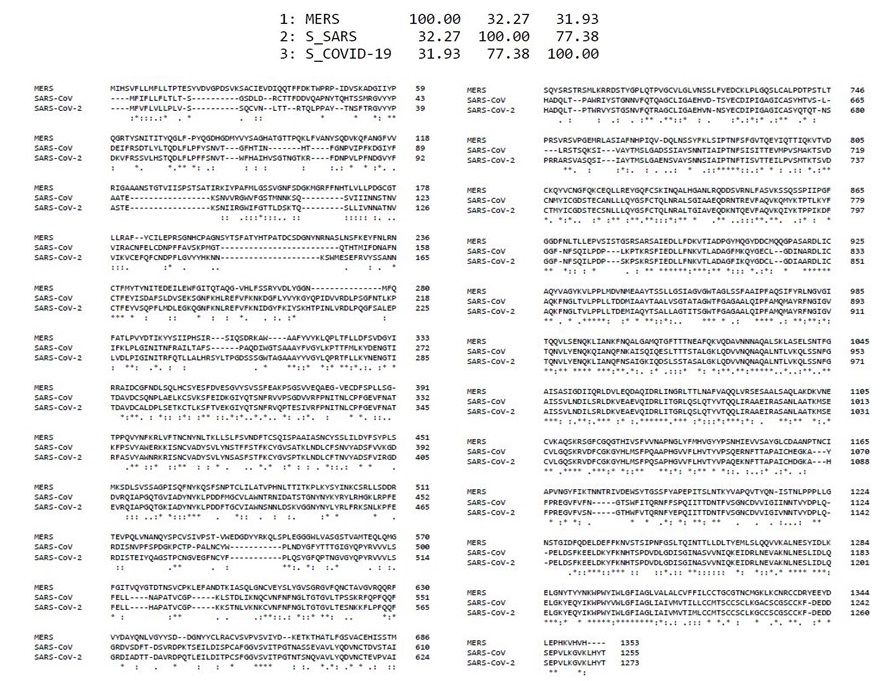

### Supplementary Fig. 3

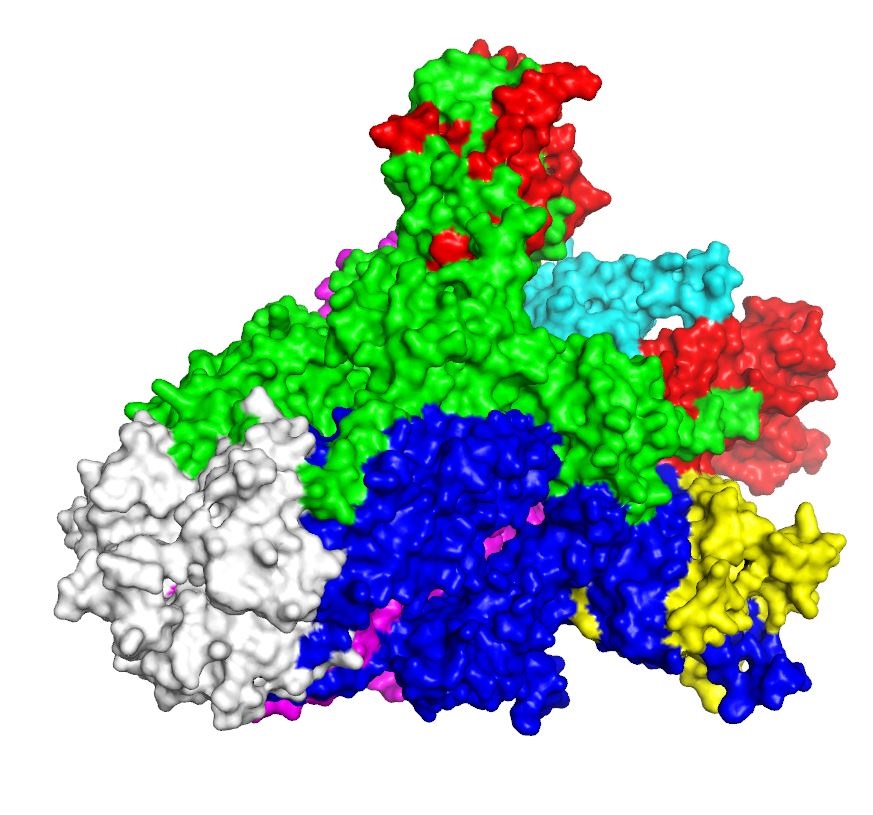

### Supplementary Fig. 4

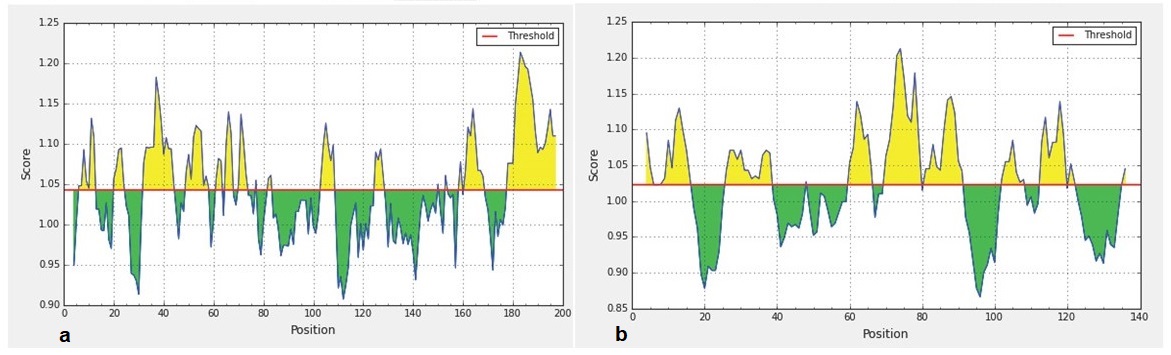

### Supplementary Fig. 5

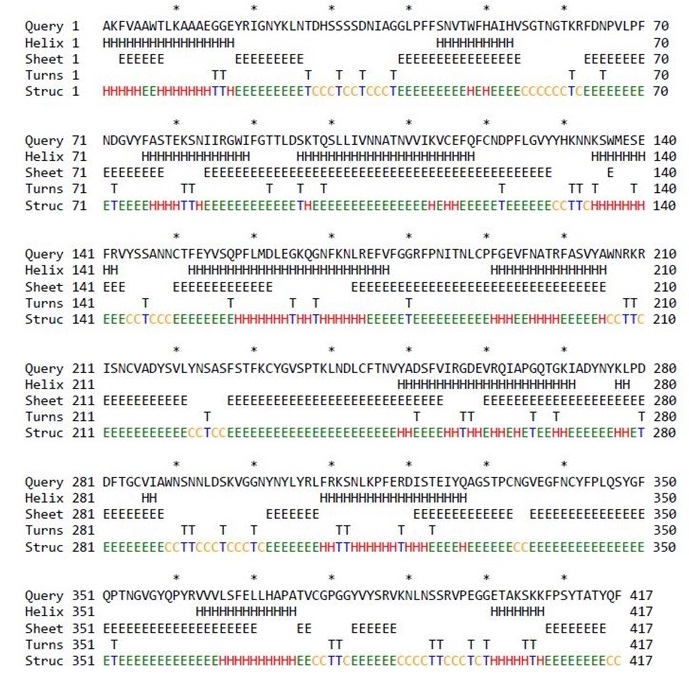

### Supplementary Fig. 6

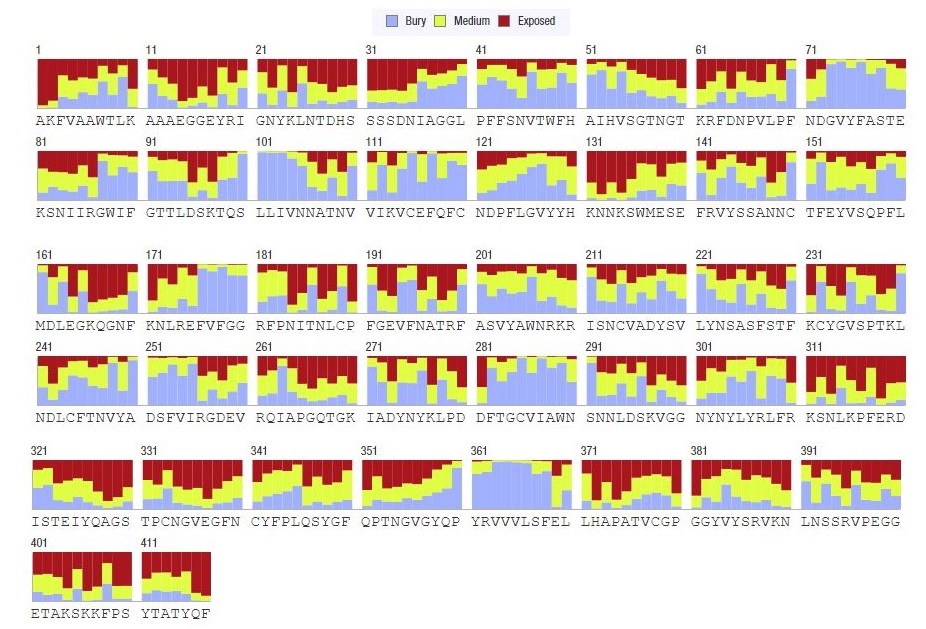

### Supplementary Fig. 7

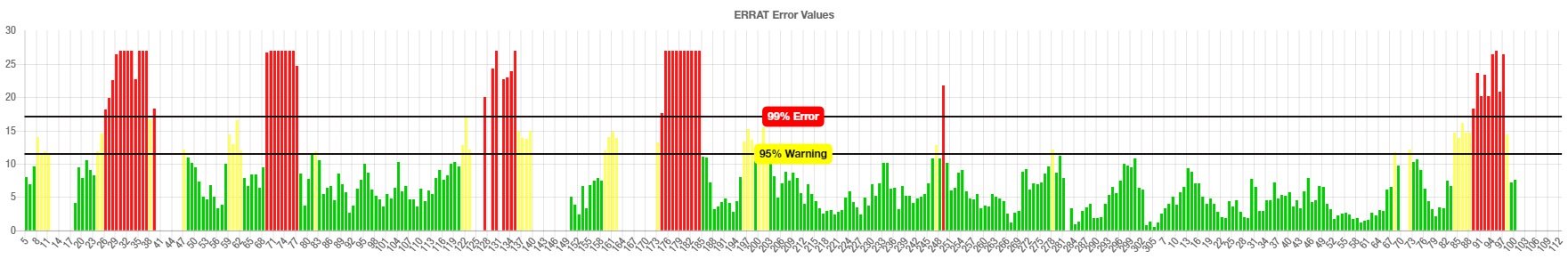
