## Supplementary Figure Legends for "Epitope-based chimeric peptide vaccine design against S, M and E proteins of SARS-CoV-2 etiologic agent of global pandemic COVID-19: an *in silico* approach"

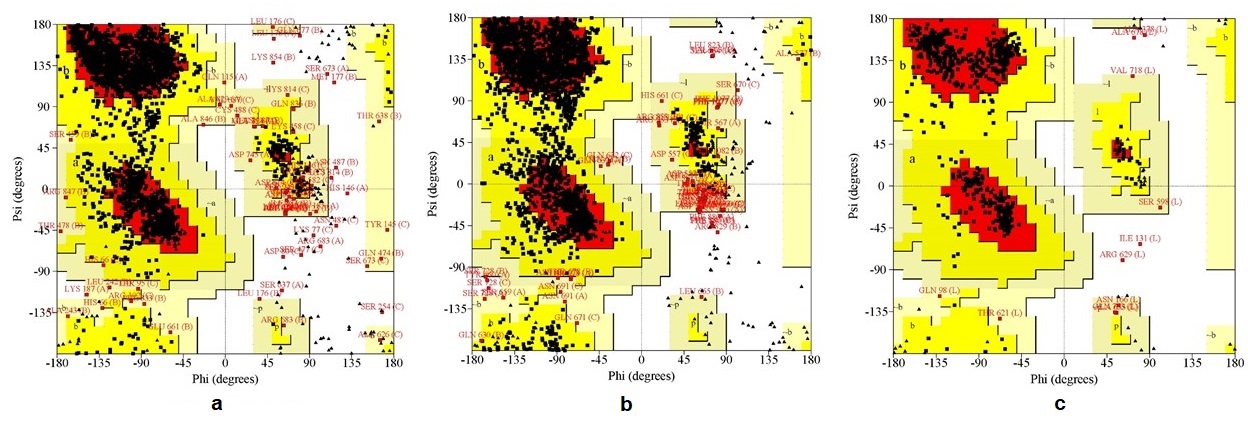

**Supplementary Fig. 1 The Ramachandran plots derived from homology modeling.** The three plots (a), (b) and (c) respectively illustrates the spike (S) proteins of SARS-CoV-2, SARS-CoV and MERS-CoV of Ramachandran outputs using PROCHECK web server. Most favored regions in the plots of are shown in red, additional allowed regions are shown in yellow, generously allowed regions are shown light brown, and disallowed regions are shown in white. After refinement, in SARS-CoV-2 S protein 85.1% and 12.7% amino acid residues were found in favored and allowed regions, respectively. The SARS-CoV S protein however had 78.4% and10.2% residues in favored and allowed regions, respectively, and 88.1% and 19.6% residues belonged to favored and allowed regions, respectively in MERS-CoV S protein.

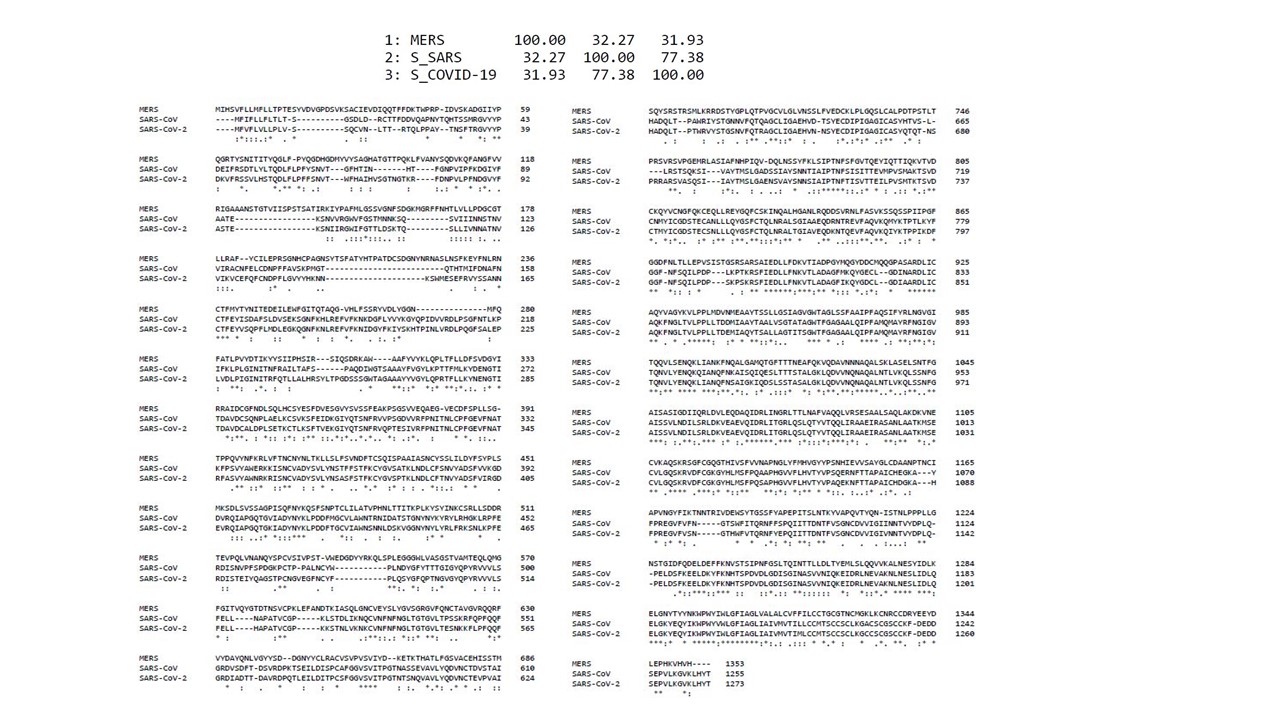

**Supplementary Fig. 2 Linear sequence alignment of the spike (S) proteins of the SARS-CoV-2, SARS-CoV and MERS-CoV.** Using ClustalW multiple sequence alignment tool (version 1.2.4) we found that the S protein of SARS-CoV-2 shares 77.38% and 31.93% sequence identity with the S proteins of the SARS-CoV and MERS-CoV, respectively.

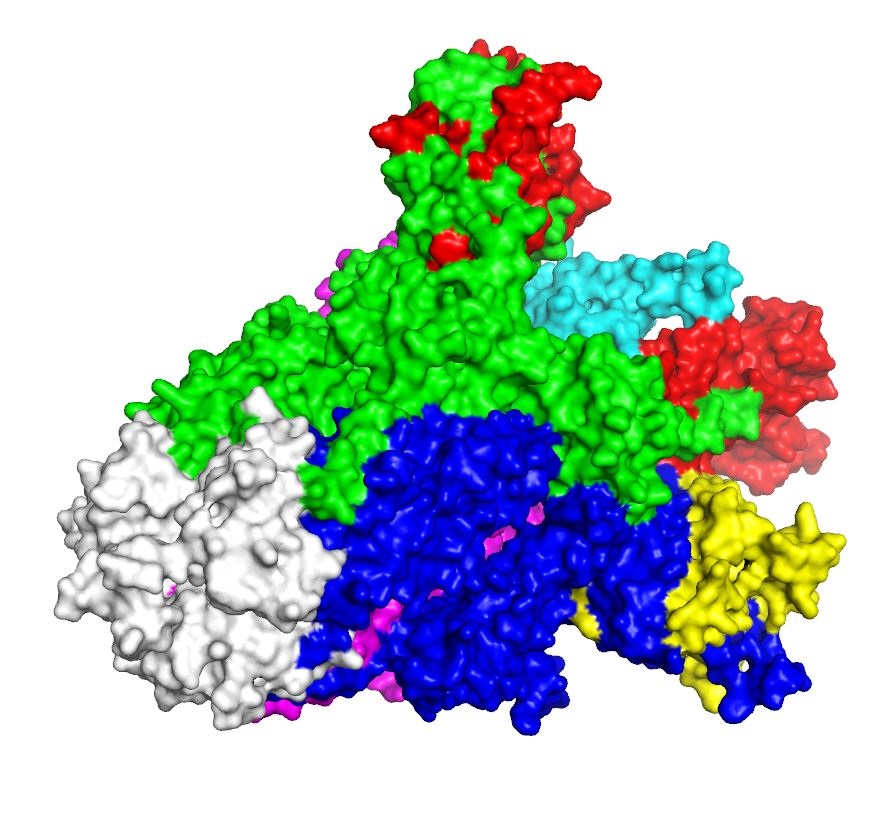

**Supplementary Fig. 3** **Three-dimensional (3D) structure of the spike (S) proteins of SARS-CoV-2 (surface view).** The red, cyan, and yellow colored regions represent the potential antigenic domains predicted by the IEDB analysis resource Elipro analysis whereas the gray colored region represents the transmembrane domain of S protein.

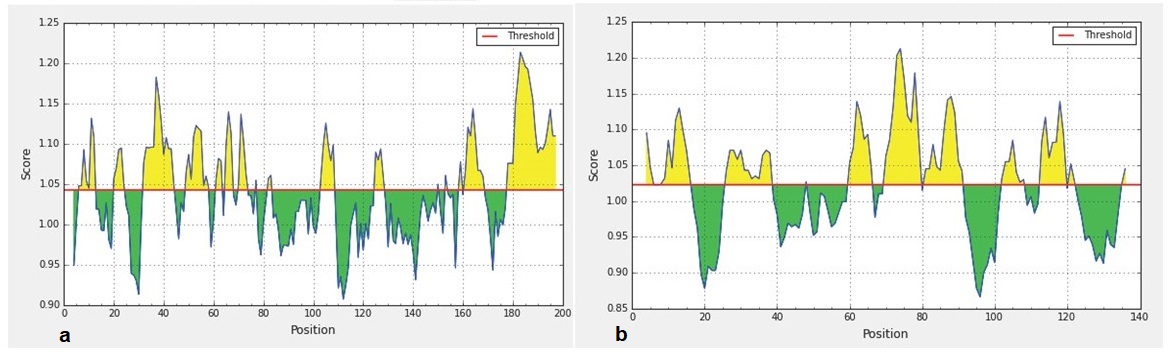

**Supplementary Fig. 4 Predicted B-cell epitopes using Kolaskar and Tongaonkar antigenicity profiling in IEDB-analysis resource web-based repository.** Yellow areas above threshold (red line) are proposed to be a part of B cell epitopes in (a) RBD and (b) NTD regions of S protein of the SARS-CoV-2. While green areas are not.

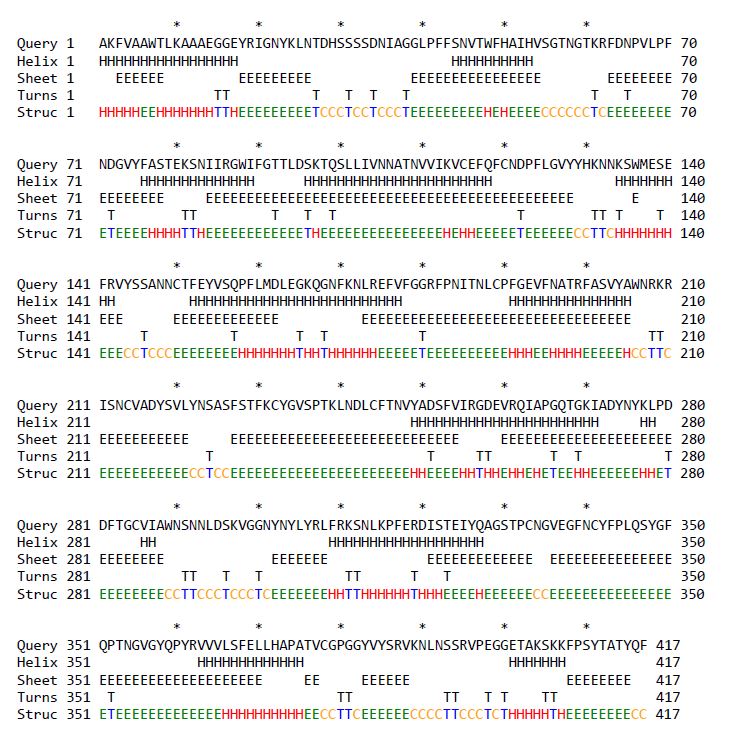

**Supplementary Fig. 5 Predicted secondary structure of CoV-RMEN using CFSSP:Chou and Fasman secondary structure prediction server.**

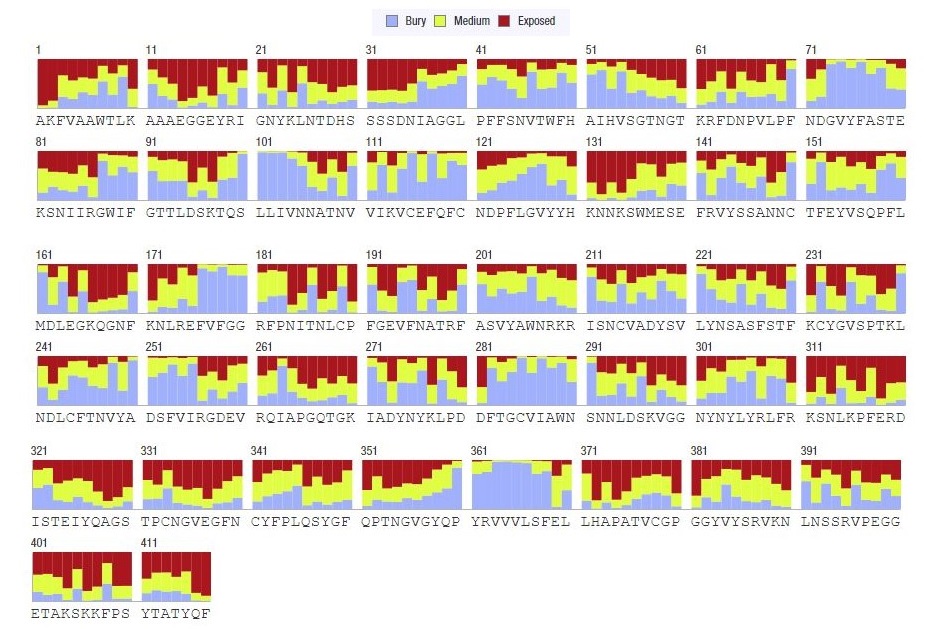

**Supplementary Fig. 6 Graphical representation of solvent accessibility of CoV-RMEN vaccine candidate sequence.**

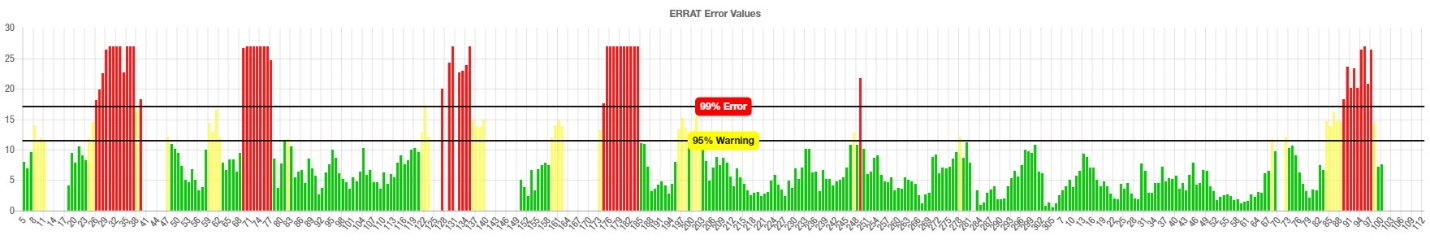

**Supplementary Fig.7 Graphical representation of the 3D structure validation of the CoV-RMEN vaccine candidate using ERRAT on-line server.**
