## Supplementary Table 1 for "Epitope-based chimeric peptide vaccine design against S, M and E proteins of SARS-CoV-2 etiologic agent of global pandemic COVID-19: an *in silico* approach"

**Supplementary Table 1:** Predicted B-cell epitopes in RBD and NTD regions of S glycoprotein, envelop (E) and membrane (M) proteins of the SARS-CoV-2 through BepiPred-2.0 sequential B-Cell epitope predictor. The epitopes highlighted in green color were considered to be promising vaccine candidates against B-cells of SARS-CoV-2.

| **No.** | **Protein** | **Start** | **End** | **Peptide** | **Length** |
| --- | --- | --- | --- | --- | --- |
| 1 | **RBD region** | 14 | 15 | VF | 2 |
| 2 |  | 17 | 22 | ATRFAS | 6 |
| 3 |  | 24 | 36 | YAWNRKRISNCVA | 13 |
| 4 |  | 45 | 51 | ASFSTFK | 7 |
| 5 |  | 55 | 55 | V | 1 |
| 6 |  | 75 | 100 | IRGDEVRQIAPGQTGKIADYNYKLPD | 26 |
| 7 |  | 113 | 158 | NLDSKVGGNYNYLYRLFRKSNLKPFERDISTEIYQAGSTPCNGVEG | 46 |
| 8 |  | 166 | 179 | QSYGFQPTNGVGYQ | 14 |
| 1 | **NTD region** | 17 | 26 | GTNGTKRFDN | 10 |
| 2 |  | 55 | 58 | LDSK | 4 |
| 3 |  | 91 | 100 | HKNNKSWMES | 10 |
| 4 |  | 106 | 107 | SS | 2 |
| 5 |  | 109 | 109 | N | 1 |
| 6 |  | 117 | 136 | SQPFLMDLEGKQGNFKNLRE | 20 |
| 1 | **E protein** | 6 | 9 | SEET | 4 |
| 2 |  | 57 | 71 | YVYSRVKNLNSSRVP | 15 |
| 1 | **M protein** | 5 | 20 | NGTITVEELKKLLEQW | 16 |
| 2 |  | 40 | 41 | AN | 2 |
| 3 |  | 132 | 137 | PLLESE | 6 |
| 4 |  | 161 | 163 | IKD | 3 |
| 5 |  | 180 | 191 | KLGASQRVAGDS | 12 |
| 6 |  | 199 | 218 | YRIGNYKLNTDHSSSSDNIA | 20 |
