## Supplementary Table 2 for "Epitope-based chimeric peptide vaccine design against S, M and E proteins of SARS-CoV-2 etiologic agent of global pandemic COVID-19: an *in silico* approach"

**Supplementary Table 2:** Predicted B-cell epitopes in RBD and NTD regions of S glycoprotein, envelop (EBE) and membrane (MBE) proteins of the SARS-CoV-2 through Kolaskar and Tongaonkar antigenicity profiling.

| **No.** | **Start** | **End** | **Peptide** | **Average score** |
| --- | --- | --- | --- | --- |
| **RBD** (Average: 1.042 Minimum: 0.907 Maximum: 1.214) | | | | |
| 1 | 6 | 12 | TNLCPFG | 7 |
| 2 | 32 | 44 | SNCVADYSVLYNS | 13 |
| 3 | 49 | 58 | TFKCYGVSPT | 10 |
| 4 | 103 | 108 | TGCVIA | 6 |
| 5 | 161 | 168 | CYFPLQSY | 8 |
| **NTD** (Average: 1.023 Minimum: 0.866 Maximum: 1.213) | | | | |
| 1 | 8 | 16 | TWFHAIHVS | 9 |
| 2 | 26 | 38 | NPVLPFNDGVYFA | 13 |
| 3 | 60 | 66 | QSLLIVN | 7 |
| 4 | 70 | 79 | NVVIKVCEFQ | 10 |
| 5 | 81 | 91 | CNDPFLGVYYH | 11 |
| 6 | 102 | 108 | FRVYSSA | 7 |
| 7 | 113 | 119 | FEYVSQP | 7 |
| **MBE** (Average: 0.980 Minimum: 0.953 Maximum: 1.002) | | | | |
| **EBE** (Average: 1.032 Minimum: 0.947 Maximum: 1.129) | | | | |
