## Supplementary Table 3 for "Epitope-based chimeric peptide vaccine design against S, M and E proteins of SARS-CoV-2 etiologic agent of global pandemic COVID-19: an *in silico* approach"

**Supplementary Table 3:** Predicted B-cell epitopes in RBD and NTD regions of S glycoprotein of the SARS-CoV-2 through ABCPred-2.0 B-Cell epitope predictor.

| **Rank** | **Sequence** | **Start position** | **Score** |
| --- | --- | --- | --- |
| **RBD domain (Average Score :0.775)** | | | |
| 1 | GSTPCNGVEGFNCYFP | 149 | 0.91 |
| 2 | LQSYGFQPTNGVGYQP | 165 | 0.90 |
| 3 | TEIYQAGSTPCNGVEG | 143 | 0.89 |
| 4 | FERDISTEIYQAGSTP | 137 | 0.86 |
| 5 | EVRQIAPGQTGKIADY | 79 | 0.85 |
| 5 | CFTNVYADSFVIRGDE | 64 | 0.85 |
| 6 | TGKIADYNYKLPDDFT | 88 | 0.84 |
| 7 | FPNITNLCPFGEVFNA | 2 | 0.82 |
| 8 | FASVYAWNRKRISNCV | 20 | 0.81 |
| 9 | NCVADYSVLYNSASFS | 33 | 0.79 |
| 10 | NGVGYQPYRVVVLSFE | 174 | 0.77 |
| 11 | FSTFKCYGVSPTKLND | 47 | 0.76 |
| 12 | SVLYNSASFSTFKCYG | 39 | 0.74 |
| 13 | EGFNCYFPLQSYGFQP | 157 | 0.73 |
| 14 | YKLPDDFTGCVIAWNS | 96 | 0.71 |
| 14 | VGGNYNYLYRLFRKSN | 118 | 0.71 |
| 14 | TGCVIAWNSNNLDSKV | 103 | 0.71 |
| 15 | FVIRGDEVRQIAPGQT | 73 | 0.69 |
| 16 | VVLSFELLHAPATVCG | 184 | 0.66 |
| 17 | RKSNLKPFERDISTEI | 130 | 0.65 |
| 18 | PTKLNDLCFTNVYADS | 57 | 0.62 |
| **NTD (Average : 0.733)** | | | |
| 1 | SWMESEFRVYSSANNC | 96 | 0.86 |
| 2 | KSNIIRGWIFGTTLDS | 42 | 0.84 |
| 3 | VSGTNGTKRFDNPVLP | 15 | 0.82 |
| 4 | TNVVIKVCEFQFCNDP | 69 | 0.80 |
| 5 | VYSSANNCTFEYVSQP | 104 | 0.79 |
| 6 | GTTLDSKTQSLLIVNN | 52 | 0.78 |
| 7 | CNDPFLGVYYHKNNKS | 81 | 0.73 |
| 8 | DGVYFASTEKSNIIRG | 33 | 0.72 |
| 9 | PFLMDLEGKQGNFKNL | 119 | 0.65 |
| 10 | PVLPFNDGVYFASTEK | 27 | 0.63 |
| 10 | TKRFDNPVLPFNDGVY | 21 | 0.63 |
| 11 | LIVNNATNVVIKVCEF | 63 | 0.55 |
