## Supplementary Table 4 for "Epitope-based chimeric peptide vaccine design against S, M and E proteins of SARS-CoV-2 etiologic agent of global pandemic COVID-19: an *in silico* approach"

**Supplementary Table 4:** IFN-γ inducing epitopes predicted by IFNepitope program

| **Serial No.** | **Start-End** | **Sequence** | **Method** | **Result** | **Score** |
| --- | --- | --- | --- | --- | --- |
| **NTD (average score :0.312)** | | | | | |
| 1 | 34-49 | VYFASTEKSNIIRGW | SVM | POSITIVE | 0.810945 |
| 2 | 35-50 | YFASTEKSNIIRGWI | SVM | POSITIVE | 0.78967 |
| 3 | 36-51 | FASTEKSNIIRGWIF | SVM | POSITIVE | 0.642475 |
| 4 | 33-48 | GVYFASTEKSNIIRG | SVM | POSITIVE | 0.637418 |
| 5 | 37-52 | ASTEKSNIIRGWIFG | SVM | POSITIVE | 0.502773 |
| 6 | 38-53 | STEKSNIIRGWIFGT | SVM | POSITIVE | 0.502331 |
| 7 | 39-54 | TEKSNIIRGWIFGTT | SVM | POSITIVE | 0.481011 |
| 8 | 82-97 | DPFLGVYYHKNNKSW | SVM | POSITIVE | 0.4484 |
| 9 | 112-127 | FEYVSQPFLMDLEGK | SVM | POSITIVE | 0.413322 |
| 10 | 40-55 | EKSNIIRGWIFGTTL | SVM | POSITIVE | 0.407048 |
| 11 | 77-92 | FQFCNDPFLGVYYHK | SVM | POSITIVE | 0.35935 |
| 12 | 76-91 | EFQFCNDPFLGVYYH | SVM | POSITIVE | 0.338283 |
| 13 | 32-47 | DGVYFASTEKSNIIR | SVM | POSITIVE | 0.305856 |
| 14 | 83-98 | PFLGVYYHKNNKSWM | SVM | POSITIVE | 0.301977 |
| 15 | 116-131 | SQPFLMDLEGKQGNF | SVM | POSITIVE | 0.2747 |
| 16 | 84-99 | FLGVYYHKNNKSWME | SVM | POSITIVE | 0.264761 |
| 17 | 81-96 | NDPFLGVYYHKNNKS | SVM | POSITIVE | 0.254982 |
| 18 | 113-128 | EYVSQPFLMDLEGKQ | SVM | POSITIVE | 0.25338 |
| 19 | 79-94 | FCNDPFLGVYYHKNN | SVM | POSITIVE | 0.239539 |
| 20 | 42-57 | SNIIRGWIFGTTLDS | SVM | POSITIVE | 0.226778 |
| 21 | 114-129 | YVSQPFLMDLEGKQG | SVM | POSITIVE | 0.210321 |
| 22 | 115-130 | VSQPFLMDLEGKQGN | SVM | POSITIVE | 0.198977 |
| 23 | 85-100 | LGVYYHKNNKSWMES | SVM | POSITIVE | 0.19694 |
| 24 | Jan-15 | PFFSNVTWFHAIHVS | SVM | POSITIVE | 0.171456 |
| 25 | 78-93 | QFCNDPFLGVYYHKN | SVM | POSITIVE | 0.169759 |
| 26 | 111-126 | TFEYVSQPFLMDLEG | SVM | POSITIVE | 0.15245 |
| 27 | 86-101 | GVYYHKNNKSWMESE | SVM | POSITIVE | 0.150871 |
| 28 | 41-56 | KSNIIRGWIFGTTLD | SVM | POSITIVE | 0.10791 |
| 29 | 43-58 | NIIRGWIFGTTLDSK | SVM | POSITIVE | 0.10336 |
| 30 | 80-95 | CNDPFLGVYYHKNNK | SVM | POSITIVE | 0.098891 |
| 31 | 71-86 | VIKVCEFQFCNDPFL | SVM | POSITIVE | 0.094081 |
| 32 | 29-44 | PFNDGVYFASTEKSN | SVM | POSITIVE | 0.087175 |
| 33 | 117-132 | QPFLMDLEGKQGNFK | SVM | POSITIVE | 0.085353 |
| **RBD (Average score :0.255)** | | | | |  |
| 1 | 135-150 | PFERDISTEIYQAGS | SVM | POSITIVE | 0.624815 |
| 2 | 136-151 | FERDISTEIYQAGST | SVM | POSITIVE | 0.572874 |
| 3 | 180-195 | YRVVVLSFELLHAPA | SVM | POSITIVE | 0.571759 |
| 4 | 133-148 | LKPFERDISTEIYQA | SVM | POSITIVE | 0.553997 |
| 5 | 18-33 | RFASVYAWNRKRISN | SVM | POSITIVE | 0.531663 |
| 6 | 138-153 | RDISTEIYQAGSTPC | SVM | POSITIVE | 0.525355 |
| 7 | 19-34 | FASVYAWNRKRISNC | SVM | POSITIVE | 0.522057 |
| 8 | 17-32 | TRFASVYAWNRKRIS | SVM | POSITIVE | 0.521612 |
| 9 | 179-194 | PYRVVVLSFELLHAP | SVM | POSITIVE | 0.520905 |
| 10 | 134-149 | KPFERDISTEIYQAG | SVM | POSITIVE | 0.514967 |
| 11 | 176-191 | GYQPYRVVVLSFELL | SVM | POSITIVE | 0.500094 |
| 12 | 137-152 | ERDISTEIYQAGSTP | SVM | POSITIVE | 0.490169 |
| 13 | 139-154 | DISTEIYQAGSTPCN | SVM | POSITIVE | 0.480013 |
| 14 | 20-35 | ASVYAWNRKRISNCV | SVM | POSITIVE | 0.474759 |
| 15 | 22-37 | VYAWNRKRISNCVAD | SVM | POSITIVE | 0.438636 |
| 16 | 175-190 | VGYQPYRVVVLSFEL | SVM | POSITIVE | 0.394901 |
| 17 | 62-77 | LCFTNVYADSFVIRG | SVM | POSITIVE | 0.366524 |
| 18 | 23-38 | YAWNRKRISNCVADY | SVM | POSITIVE | 0.35601 |
| 19 | 21-36 | SVYAWNRKRISNCVA | SVM | POSITIVE | 0.350959 |
| 20 | 178-193 | QPYRVVVLSFELLHA | SVM | POSITIVE | 0.345882 |
| 21 | 181-196 | RVVVLSFELLHAPAT | SVM | POSITIVE | 0.324276 |
| 22 | 177-192 | YQPYRVVVLSFELLH | SVM | POSITIVE | 0.318673 |
| 23 | 61-76 | DLCFTNVYADSFVIR | SVM | POSITIVE | 0.282876 |
| 24 | 140-155 | ISTEIYQAGSTPCNG | SVM | POSITIVE | 0.256374 |
| 25 | 16-31 | ATRFASVYAWNRKRI | SVM | POSITIVE | 0.243891 |
| 26 | 79-94 | VRQIAPGQTGKIADY | SVM | POSITIVE | 0.238696 |
| 27 | 143-158 | EIYQAGSTPCNGVEG | SVM | POSITIVE | 0.233029 |
| 28 | 182-197 | VVVLSFELLHAPATV | SVM | POSITIVE | 0.229693 |
| 29 | 81-96 | QIAPGQTGKIADYNY | SVM | POSITIVE | 0.22687 |
| 30 | 141-156 | STEIYQAGSTPCNGV | SVM | POSITIVE | 0.193904 |
| 31 | 183-198 | VVLSFELLHAPATVC | SVM | POSITIVE | 0.176479 |
| 32 | 15-30 | NATRFASVYAWNRKR | SVM | POSITIVE | 0.162741 |
| 33 | 27-42 | RKRISNCVADYSVLY | SVM | POSITIVE | 0.152176 |
| 34 | 80-95 | RQIAPGQTGKIADYN | SVM | POSITIVE | 0.143402 |
| 35 | 174-189 | GVGYQPYRVVVLSFE | SVM | POSITIVE | 0.12467 |
| 36 | 26-41 | NRKRISNCVADYSVL | SVM | POSITIVE | 0.120148 |
| 37 | 116-131 | KVGGNYNYLYRLFRK | SVM | POSITIVE | 0.114659 |
| 38 | 63-78 | CFTNVYADSFVIRGD | SVM | POSITIVE | 0.112158 |
| 39 | 24-39 | AWNRKRISNCVADYS | SVM | POSITIVE | 0.107934 |
| 40 | 38-53 | SVLYNSASFSTFKCY | SVM | POSITIVE | 0.106128 |
| 41 | 39-54 | VLYNSASFSTFKCYG | SVM | POSITIVE | 0.088337 |
| 42 | 165-180 | QSYGFQPTNGVGYQP | SVM | POSITIVE | 0.076122 |
| 43 | 117-132 | VGGNYNYLYRLFRKS | SVM | POSITIVE | 0.076019 |
| 44 | 82-97 | IAPGQTGKIADYNYK | SVM | POSITIVE | 0.068857 |
| 45 | Sep-24 | PFGEVFNATRFASVY | SVM | POSITIVE | 0.064041 |
| 46 | 118-133 | GGNYNYLYRLFRKSN | SVM | POSITIVE | 0.06335 |
| 47 | 13-28 | VFNATRFASVYAWNR | SVM | POSITIVE | 0.05905 |
| 48 | Nov-26 | GEVFNATRFASVYAW | SVM | POSITIVE | 0.058894 |
| 49 | Dec-27 | EVFNATRFASVYAWN | SVM | POSITIVE | 0.043785 |
| 50 | 64-79 | FTNVYADSFVIRGDE | SVM | POSITIVE | 0.043701 |
| 51 | 132-147 | NLKPFERDISTEIYQ | SVM | POSITIVE | 0.030851 |
| 52 | 142-157 | TEIYQAGSTPCNGVE | SVM | POSITIVE | 0.024497 |
| 53 | 28-43 | KRISNCVADYSVLYN | SVM | POSITIVE | 0.016069 |
| 54 | 156-171 | EGFNCYFPLQSYGFQ | SVM | POSITIVE | 0.011051 |
| 55 | 184-199 | VLSFELLHAPATVCG | SVM | POSITIVE | 0.009809 |
| 56 | 30-45 | ISNCVADYSVLYNSA | SVM | POSITIVE | 6.87E-05 |
| MBE POSITIVE 0.97925313 | | | | | |
