## Supplementary Table 5 for "Epitope-based chimeric peptide vaccine design against S, M and E proteins of SARS-CoV-2 etiologic agent of global pandemic COVID-19: an *in silico* approach"

Supplementary Table 5: NCBI partial or complete genome sequences accession no and their respective country.

| **Accession no.** | **Country** |
| --- | --- |
| MN908947 | China |
| MN970003 | Thailand |
| MN970004 | Thailand |
| MN938384 | China |
| MN938385 | China |
| MN938386 | China |
| MN938387 | China |
| MN938388 | China |
| MN938389 | China |
| MN938390 | China |
| MN975262 | China |
| MN975263 | China |
| MN975264 | China |
| MN975265 | China |
| MN975266 | China |
| MN975267 | China |
| MN975268 | China |
| MN985325 | USA |
| MN988713 | USA |
| MN994467 | USA |
| MN994468 | USA |
| MN997409 | USA |
| MN988668 | China |
| MN988669 | China |
| MN996527 | China |
| MN996528 | China |
| MN996529 | China |
| MN996530 | China |
| MN996531 | China |
| MT007544 | Australia |
| MT008022 | Italy |
| MT008023 | Italy |
| LR757995 | China |
| LR757996 | China |
| LR757997 | China |
| LR757998 | China |
| MT019529 | China |
| MT019530 | China |
| MT019531 | China |
| MT019532 | China |
| MT019533 | China |
| MT020781 | Finland |
| MT020880 | USA |
| MT020881 | USA |
| MT027062 | USA |
| MT027063 | USA |
| MT027064 | USA |
| LC522350 | Philippines |
| MT039873 | China |
| MT039887 | USA |
| MT039888 | USA |
| MT039890 | South Korea |
| MT042773 | China |
| MT042774 | China |
| MT042775 | China |
| MT042776 | China |
| MT042777 | China |
| MT042778 | China |
| MT044257 | USA |
| MT044258 | USA |
| MT050414 | Australia |
| MT050415 | Australia |
| MT050416 | Australia |
| MT050417 | Australia |
| MT049951 | China |
| LC523807 | Philippines |
| LC523808 | Philippines |
| LC523809 | Malaysia |
| MT066157 | Malaysia |
| MT066158 | Malaysia |
| MT066159 | Taiwan |
| MT066175 | Taiwan |
| MT066176 | Belgium |
| MT072667 | Belgium |
| MT072668 | Nepal |
| MT072688 | China |
| MT081059 | China |
| MT081060 | China |
| MT081061 | China |
| MT081062 | China |
| MT081063 | China |
| MT081064 | China |
| MT081065 | China |
| MT081066 | China |
| MT081067 | China |
| MT081068 | Sweden |
| MT093571 | China |
| MT093631 | USA |
| MT106052 | USA |
| MT106053 | USA |
| MT106054 | Australia |
| MT111895 | Australia |
| MT111896 | USA |
| MT118835 | China |
| MT123290 | China |
| MT123291 | China |
| MT123292 | China |
| MT123293 | Japan |
| LC528232 | Japan |
| LC528233 | Brazil |
| MT126808 | Viet Nam |
| MT127113 | Viet Nam |
| MT127114 | Viet Nam |
| MT127115 | Viet Nam |
| MT127116 | China |
| MT135041 | China |
| MT135042 | China |
| MT135043 | China |
| MT135044 | USA |
| MT152824 | Iran |
| MT152900 | India |
| MT012098 | India |
| MT050493 | China |
| MT121215 | USA |
| MT159705 | USA |
| MT159706 | USA |
| MT159707 | USA |
| MT159708 | USA |
| MT159709 | USA |
| MT159710 | USA |
| MT159711 | USA |
| MT159712 | USA |
| MT159713 | USA |
| MT159714 | USA |
| MT159715 | USA |
| MT159716 | USA |
| MT159717 | USA |
| MT159718 | USA |
| MT159719 | USA |
| MT159720 | USA |
| MT159721 | USA |
| MT159722 | Nigeria |
| MT159778 | Italy |
| MT066156 | Italy |
| MT161607 | Iran |
| MT163712 | India |
| MT163714 | India |
| MT163715 | USA |
| MT163716 | USA |
| MT163717 | USA |
| MT163718 | USA |
| MT163719 | USA |
| MT163720 | USA |
| MT163721 | Iran |
| MT163737 | Iran |
| MT163738 | Japan |
| LC529905 | USA |
| MT184907 | USA |
| MT184908 | USA |
| MT184909 | USA |
| MT184910 | USA |
| MT184911 | USA |
| MT184912 | USA |
| MT184913 | Iran |
| MT186676 | Iran |
| MT186677 | Iran |
| MT186678 | Iran |
| MT186679 | Iran |
| MT186680 | Iran |
| MT186681 | Iran |
| MT186682 | Italy |
| MT187977 | USA |
| MT188339 | USA |
| MT188340 | USA |
| MT188341 | Italy |
| MT192758 | Taiwan |
| MT192759 | USA |
| MT192765 | Viet Nam |
| MT192772 | Viet Nam |
| MT192773 | Spain |
| MT198651 | Spain |
| MT198652 | Spain |
| MT198653 | China |
| MT226610 | Iran |
| MT232869 | Iran |
| MT232870 | Iran |
| MT232871 | Iran |
| MT232872 | Spain |
| MT233519 | Spain |
| MT233520 | Spain |
| MT233521 | Spain |
| MT233522 | Spain |
| MT233523 | Pakistan |
| MT240479 | USA |
| MT246449 | USA |
| MT246450 | USA |
| MT246451 | USA |
| MT246452 | USA |
| MT246453 | USA |
| MT246454 | USA |
| MT246455 | USA |
| MT246456 | USA |
| MT246457 | USA |
| MT246458 | USA |
| MT246459 | USA |
| MT246460 | USA |
| MT246461 | USA |
| MT246462 | USA |
| MT246463 | USA |
| MT246464 | USA |
| MT246465 | USA |
| MT246466 | USA |
| MT246467 | USA |
| MT246468 | USA |
| MT246469 | USA |
| MT246470 | USA |
| MT246471 | USA |
| MT246472 | USA |
| MT246473 | USA |
| MT246474 | USA |
| MT246475 | USA |
| MT246476 | USA |
| MT246477 | USA |
| MT246478 | USA |
| MT246479 | USA |
| MT246480 | USA |
| MT246481 | USA |
| MT246482 | USA |
| MT246483 | USA |
| MT246484 | USA |
| MT246485 | USA |
| MT246486 | USA |
| MT246487 | USA |
| MT246488 | USA |
| MT246489 | USA |
| MT246490 | USA |
| MT251972 | USA |
| MT251973 | USA |
| MT251974 | USA |
| MT251975 | USA |
| MT251976 | USA |
| MT251977 | USA |
| MT251978 | USA |
| MT251979 | USA |
| MT251980 | China |
| MT253696 | China |
| MT253697 | China |
| MT253698 | China |
| MT253699 | China |
| MT253700 | China |
| MT253701 | China |
| MT253702 | China |
| MT253703 | China |
| MT253704 | China |
| MT253705 | China |
| MT253706 | China |
| MT253707 | China |
| MT253708 | China |
| MT253709 | China |
| MT253710 | China |
